# Designing biochemical circuits with tree search

**DOI:** 10.1101/2025.01.27.635147

**Authors:** Pranav S. Bhamidipati, Matthew Thomson

## Abstract

Discovering biochemical circuits that exhibit a desired behavior is an outstanding problem in biological engineering. The traditional approach of enumerating every possible circuit topology becomes intractable for circuits with more than four components due to combinatorial scaling of the search space. Here, we use Monte Carlo Tree Search (MCTS), a reinforcement learning (RL) algorithm, to optimize circuit topology for a target phenotype by approaching circuit design as a sequence of assembly decisions. Our RL-based design framework, which we call CircuiTree, efficiently and comprehensively finds robust designs for three-component oscillators by prioritizing sparsity. CircuiTree can also infer candidate network motifs from its search results, producing similar results to enumeration. Using parallel MCTS, we scale this workflow up to five components and find that highly fault-tolerant designs use a novel strategy, which we call motif multiplexing. Multiplexed circuits contain many overlapping network motifs that each activate in different mutational scenarios, in one case lending robustness to four out of five single-gene deletions. Overall, CircuiTree provides the first scalable computational platform for designing biochemical circuits.

## Introduction

Understanding how biochemical networks produce biological behavior is one of the central goals of systems and synthetic biology. Studying the design principles of these networks, or circuits, provides fundamental insights into their computational capabilities and enables the application of synthetic biology to problems such as cell-based therapeutics (***Lim et al. (2013***); ***Williams et al. (2020***)) and engineered microbial communities (***Jones et al. (2024***); ***Purnick and Weiss (2009***)). The topology, or qualitative architecture, of a synthetic gene circuit is a critical determinant of its behavior, and choosing a topology is the first step in engineering circuits with novel functions such as switching in response to an input (***Gardner et al. (2000***)), spontaneous oscillations (***Elowitz and Leibler (2000***)), or multistability (***Zhu et al. (2022***)). Circuit topologies with a particular desired phenotype have traditionally been discovered *de novo* using a method we call computational enumeration. Enumeration involves listing every topology in a predefined space of possibilities and simulating its behavior with an exhaustive set of random perturbations of the modeling parameters (***Ma et al. (2009***); ***Cotterell and Sharpe (2010***); ***Chau et al. (2012***); ***Schaerli et al. (2014***); ***Gerardin et al. (2019***)). This method identifies the topologies that are the most robust to parameter choice, as well as the network motifs of the phenotype, which are topological patterns overrepresented among highly robust circuits (***Alon (2007***, 2019)).

Currently, computational complexity has limited the role of automation in circuit design. Assuming 3 possible types of interactions (activation, inhibition, or no interaction), the number of distinct topologies that can be made with *N* circuit components scales as 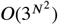, not accounting for symmetries. Because the enumeration method requires simulating each topology in this potential space with many parameter perturbations (typically ≥ 10^4^), designing a five-component circuit with this approach would require on the order of 10^15^-10^16^ simulations, a computationally intractable number. This “curse of dimensionality” constitutes the main barrier to understanding the structure-function relationship for large circuits. The search space can be narrowed by only considering combinations of smaller motifs (***Qiao et al. (2019***)), but this approach risks excluding novel, non-modular designs that appear at higher complexity (***Jiménez et al. (2017***)). For large circuits, a functional topology may be found with evolutionary optimization (***François et al. (2007***); ***François and Siggia (2010***)) or recurrent neural network-guided inference (***Shen et al. (2021***)), but these methods can be sensitive to the optimization parameters and the nature of the fitness landscape, a fact that complicates inference of network motifs and other general design principles. The ideal design framework for circuit topologies would be: (i) *Efficient*: Using relatively few samples, it returns a set of reasonably robust solutions. (ii) *Generalizable*: It generates solutions that are representative of the ground truth, enabling inference of motifs and design principles. (iii) *Scalable:* It maintains features (i) and (ii) when searching large spaces of potential designs.

Our inspiration for a circuit design approach with these features comes from algorithms developed in modern artificial intelligence to solve decision problems, which share core computational similarities with genetic circuit design. In a decision problem, or game, one starts from an initial position (the root state) and makes a sequence of decisions ending in success or failure. Algorithms for optimal game playing sample the branching tree of possible decisions in search of the best decision sequences, or paths. Large game trees, characteristic of long, complex games like chess (~ 10^43^ states) and Go (10^170^ states), can overwhelm classical exhaustive search methods (e.g., flat search, *A*^∗^, and minimax) and branch-and-bound methods (e.g., *α*-*β* pruning) (***Shannon (1950***); ***Tromp and Farnebäck (2007***)). In contrast, most modern probabilistic search algorithms use reinforcement learning (RL) to sample decision paths; for example, Monte Carlo tree search (MCTS) uses an RL decision rule that optimistically estimates the success of each possible decision (***Kocsis and Szepesvári (2006***)). MCTS has proven to be instrumental in solving large games (***Silver et al. (2016***, 2018)) due to its low memory footprint, ability to be parallelized (***Enzenberger and Müller (2010***)), balance of exploitation and exploration (***Auer et al. (2002***)), and myriad variations (***Świechowski et al. (2022***); ***Browne et al. (2012***)).

In this work, we present a computational platform that generates biological circuit designs to specification by learning exclusively from stochastic simulations. This platform, which we call CircuiTree, approaches circuit design as a sequence of assembly decisions and optimizes the circuit topology for a given phenotype by using MCTS to search the tree of possible circuit assemblies. We first used CircuiTree to search a trial space — all 3,325 connected circuits with three transcription factors — for topologies that exhibit spontaneous, sustained oscillations. As it dynamically explores the search space, CircuiTree discovers sparse oscillators before complex ones and reliably identifies the most robust topologies. Next, we demonstrate that CircuiTree accurately identifies assembly motifs, which are overrepresented search states analogous to network motifs found by enumeration. The best assembly motifs for 3-component oscillators also contain the activatorinhibitor (AI) and/or repressor (Rep) motifs, a result consistent with previous studies of 2- and 3-component oscillators. Finally, we use a parallelized implementation of CircuiTree to study mutational tolerance in 5-node circuits with ≤ 15 interactions, representing approximately 2 billion possible topologies. We discover that highly fault-tolerant five-component oscillator circuits are “multiplexed” in that they contain multiple AI and Rep motifs that are synergistically interleaved such that the oscillations are robust to complete and partial knockdowns of single genes. Our results demonstrate that CircuiTree’s RL-based framework enables the efficient, generalizable, and scalable design of biochemical circuits, which could be used to engineer and understand complex intracellular and multicellular biological systems.

## Results

### Circuit design as an assembly game

We define a circuit as a system of *k* dynamically interacting biochemical species **x** = [*x*_0_, …, *x*_*k*_] characterized by its qualitative topology *s*, a member of the set of possible topologies 𝒮, and its parameters *θ* ∈ ℝ^*n*(*s*)^, which describe the rates of biochemical reactions. We also assume the existence of a phenotype function *f* (**x**(*t*)) = *q* ∈ {0, 1} that returns a yes-or-no verdict as to whether a given simulation **x**(*t*) exhibits the desired phenotype. This function, which we conceptualize quite generally, may represent the presence of a dynamical phenotype such as adaptation or oscillation or a more general property such as multistability. Like many prior computational studies of design principles (***Ma et al. (2009***); ***Chau et al. (2012***); ***Cotterell and Sharpe (2010***); ***Schaerli et al. (2014***)), each simulation of the system is performed with a set of randomly chosen parameters, and we define robustness *Q*(*s*) as the mean phenotype score from these samples *Q*(*s*) = 𝔼_*θ*_[*q*(*s*)] = 𝔼_*θ*_ [*f* (**x**(*t*; *s, θ*))]. The most robust topology *s*^∗^ is therefore the solution to a combinatorial optimization problem. (Note that *Q* in this context should not be confused with the value function used in RL.)

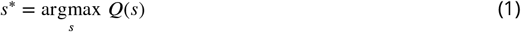

In this work, for convenience, we consider models where all topologies are governed by the same reaction parameters (i.e. *n*(*s*_*i*_) = *n*(*s*_*j*_)∀*s*_*i*_, *s*_*j*_ ∈ 𝒮), but CircuiTree does not require this assumption.

To solve Equation 1, we approach circuit topology design as a multi-step decision problem, or game (Figure 1A). Beginning with an “empty” circuit *s*_0_ with a given set of components and without interactions, each step of the game can add an activating or inhibitory (auto-)regulatory interaction. The game ends when the builder chooses to “terminate” the assembly at some topology *s*_*i*_, at which point the outcome is decided by a simulation with a win probability *Q*(*s*_*i*_). In the language of Markov decision processes (MDPs), each topology *s* represents a state of the game, and the addition of an interaction or the termination of assembly represents an action *a* that converts a state *s*_*i*_ to a different state *s*_*j*_. Each playout of the game traces a path on the tree of possible decisions *T* rooted at *s*_0_. The nodes in the tree are circuit topologies 𝒮, and each directed edge (*s*_*i*_, *s*_*j*_) ∈ *ε* represents the assembly of *s*_*j*_ from *s*_*i*_ by an action. Note that because multiple decision paths can converge on the same *s*_*j*_, *T* is technically a directed acyclic graph (DAG), but we will call it a tree for simplicity.

**Figure 1.**
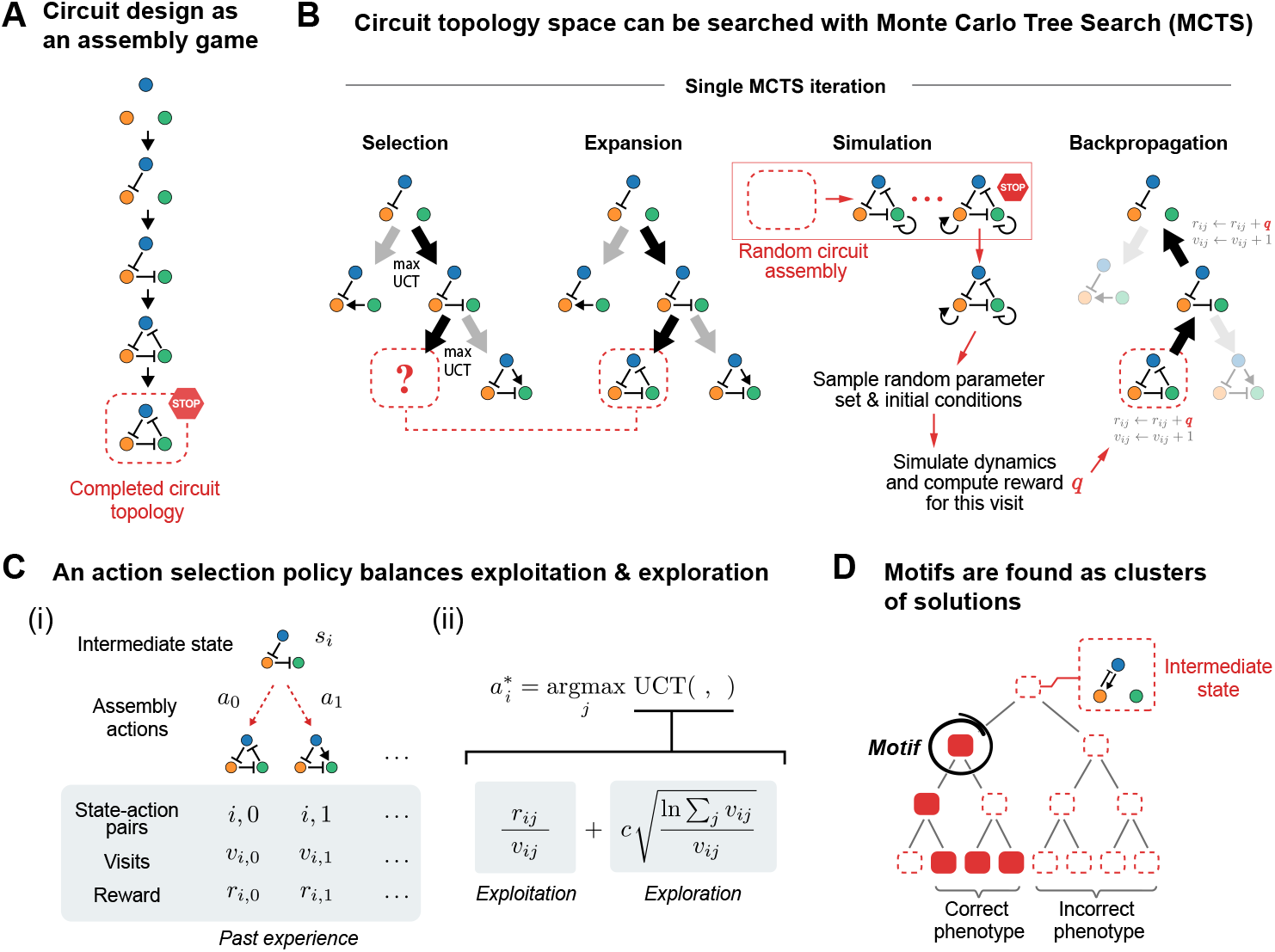
A stepwise assembly framework enables circuit topology optimization with tree search. (A) Circuit topologies are built step-by-step by adding interactions until the game is ended by taking the “terminate” action (the STOP sign). (B) Each MCTS iteration undergoes four phases: (1) Selection: The UCT criterion is used to recursively select the most promising action 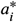 from the current state *s*_*i*_. (2) Expansion: If the edge 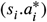 has not been sampled yet, it is added to the tree. (3) Simulation: If the circuit has not been completed yet, take random assembly moves until “terminate” is chosen. Simulate the resulting topology with random parameters and evaluate the reward *q* based on a target phenotype. (4) Backpropagation: update the history of rewards *r*_*ij*_ over past visits *v*_*ij*_ for the visited edges 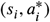. (C) The UCT policy for selecting 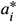 balances exploitation and exploration based on past trials. The exploration term, tuned by a hyperparameter *c*, favors actions that are under-sampled relative to the alternatives. (D) Once assembled, a motif creates an enriched subgraph in the search graph.

For large design spaces, it is computationally infeasible to estimate *Q*(*s*) for every *s* ∈ *T*. Instead, we iteratively sample from *T* using Monte Carlo tree search (MCTS), an RL algorithm commonly used in MDPs and game-playing artificial intelligences (***Silver et al. (2016***, 2017, 2018)) to bound the complexity of searching large decision trees. In these applications, tree search is used during game play (and sometimes also during training) to find the best move by simulating many hypothetical scenarios forward in time. While prior domain knowledge can be incorporated into this search process in principle, CircuiTree learns purely through simulation, similar to the general game-playing AI AlphaZero.

Each MCTS iteration starts at the root state *s*_0_ and undergoes four phases, shown in Figure 1B and described in Algorithm 1. The key innovation of MCTS occurs during the first phase (selection), when it uses the UCT criterion from RL to choose moves while balancing (i) exploitation of moves that have yielded high rewards in the past and (ii) exploration of new moves that may yield higher rewards. At each branch (Figure 1C, left), from a state *s*_*i*_, multiple actions *a*_*j*_ are available that have yielded cumulative rewards *r*_*ij*_ over *v*_*ij*_ previous visits. MCTS decides the optimal action using the decision policy 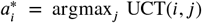 (Figure 1C, right), originally developed to solve the classic multi-armed bandit problem (***Auer et al. (2002***)).

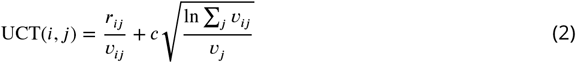

The first term of the UCT criterion estimates the mean reward of the move *s*_*i*_ → *s*_*j*_, while the second term, derived from Hoeffding’s inequality (***Kocsis and Szepesvári (2006***)), is an estimated upper bound on the sampling error. The second term ensures that a move with a historically low reward but relatively few samples (*v*_*j*_ << Σ_*j*_ *v*_*ij*_) will still be tried occasionally to account for insufficient sample size. Thus, UCT is an optimistic predictor of a move’s true mean reward. Note that the two terms encourage exploitation and exploration, respectively. (By definition, UCT(*i, j*) = +∞ if *s*_*i*_ → *s*_*j*_ has not been visited before).

Using the UCT criterion, actions are selected recursively, breaking any ties randomly, until the new state *s*_*j*_ is terminal (i.e. the assembly is finished) or has not been visited before during sampling. An unvisited *s*_*j*_ is added to *T*. Then, during the simulation phase, the outcome *q* of the trial is determined by taking random actions until the “terminate” action is chosen and simulating the resulting topology with randomly chosen reaction parameters and initial conditions. Finally, the history of rewards and visits is updated for each selected edge in *T* (backpropagation).

In addition to finding individual circuit topologies, an important goal of circuit design is to identify specific structural features, or network motifs, that lead to successful designs (***Milo et al. (2002***); ***Alon (2007***); ***Ma et al. (2009***); ***Cotterell and Sharpe (2010***); ***Shah and Sarkar (2011***); ***Chau et al. (2012***); ***Lim et al. (2013***); ***Schaerli et al. (2014***)). Similarly, in our game-playing paradigm, a motif of the assembly game is a series of moves (an assembly *strategy*) that leads to topologies with a high rate of phenotypic success, on average. Thus, when a motif is present in the space of possible designs, it creates an enriched region in *T*, which can be exploited during the tree search to rapidly discover successful designs (Figure 1D).

### Establishing a ground-truth for three-node stochastic oscillators with enumeration

Oscillations appear in diverse cellular contexts such as metabolism (***Chandra et al. (2011***)), DNA damage (***Geva-Zatorsky et al. (2006***, 2010)), the cell cycle (***Ferrell et al. (2011***)), and circadian rhythms (***Tyson et al. (1999***)), and consequently, many of their design principles have been elucidated (***Novák and Tyson (2008***)). We first benchmark CircuiTree on the well-studied problem of designing an oscillator circuit with three nodes (Figure 2A, part i). Specifically, we consider a system of symmetric transcription factors (TFs) modeled as a stochastic birth-and-death process of individual mR-NAs, TFs, and TF-response element (TF-RE) complexes, with elementary reactions of transcription, translation, degradation, binding, and unbinding (Figure S1A). Reactions and their rate parameters are summarized in Table 1; unless otherwise specified, the values used are *k*_on_ = 1.0 sec^−1^, *k*_off,1_ = 99.0 molec^−1^ sec^−1^, *k*_off,2_ = 9.9 molec^−1^ sec^−1^, *k*_unbound_ = *k*_mixed_ = 0.05 sec^−1^, *k*_act_ = 8.0 sec^−1^, *k*_inh_ = 5 × 10^−4^ sec^−1^, *k*_*p*_ = 0.167 molec^−1^ sec^−1^, and γ_*m*_ = γ_*p*_ = 0.025 molec^−1^ sec^−1^. TFs regulate transcription by binding sequentially to two REs in the regulatory region of each promoter, and cooperativity is introduced by assuming that when both sites are occupied by the same TF, the second TF-RE binding event is stronger (*K*_*D*,2_ < *K*_*D*,1_). Notably, the effective cooperativity of sequential binding can be shown to have a Hill coefficient *n*_*H*_ ≤ 2 (see Methods for a derivation of this property). For each stochastic simulation, the system was randomly initialized and stochastically simulated using the Gillespie method for *t*_max_ = 4 ×10^4^ sec = 11.1 hrs (see Methods for details of initialization). Oscillations were quantified by computing the normalized autocorrelation function (ACF) and identifying its lowest minimum value across all TFs ACF_min_, excluding the bounds, and a Boolean reward value was assigned based on a threshold value ACF_min_ (Figure 2A, part ii).

**Table 1.** Model rate parameters.

| Reaction | Rate parameter | Value in Fig. 4D & Fig. 5 |
| --- | --- | --- |
| TF-RE binding | $k_{\text{on}}$ | $1.0 \text{ sec}^{-1}$ |
| TF-RE unbinding (uncooperative) | $k_{\text{off},1}$ | $99.0 \text{ molec}^{-1} \text{ sec}^{-1}$ |
| TF-RE unbinding (cooperative) | $k_{\text{off},2}$ | $9.9 \text{ molec}^{-1} \text{ sec}^{-1}$ |
| Transcription, basal | $k_{\text{unbound}}$ | $0.05 \text{ sec}^{-1}$ |
| Transcription, 1–2 activators | $k_{\text{act}}$ | $8.0 \text{ sec}^{-1}$ |
| Transcription, 1–2 inhibitors | $k_{\text{inh}}$ | $5 \times 10^{-4} \text{ sec}^{-1}$ |
| Transcription, 1 activator and 1 inhibitor | $k_{\text{mixed}}$ | $0.05 \text{ sec}^{-1}$ |
| Translation | $k_p$ | $0.167 \text{ molec}^{-1} \text{ sec}^{-1}$ |
| Degradation, mRNA | $\gamma_m$ | $0.025 \text{ molec}^{-1} \text{ sec}^{-1}$ |
| Degradation, protein | $\gamma_p$ | $0.025 \text{ molec}^{-1} \text{ sec}^{-1}$ |

**Figure 2.**
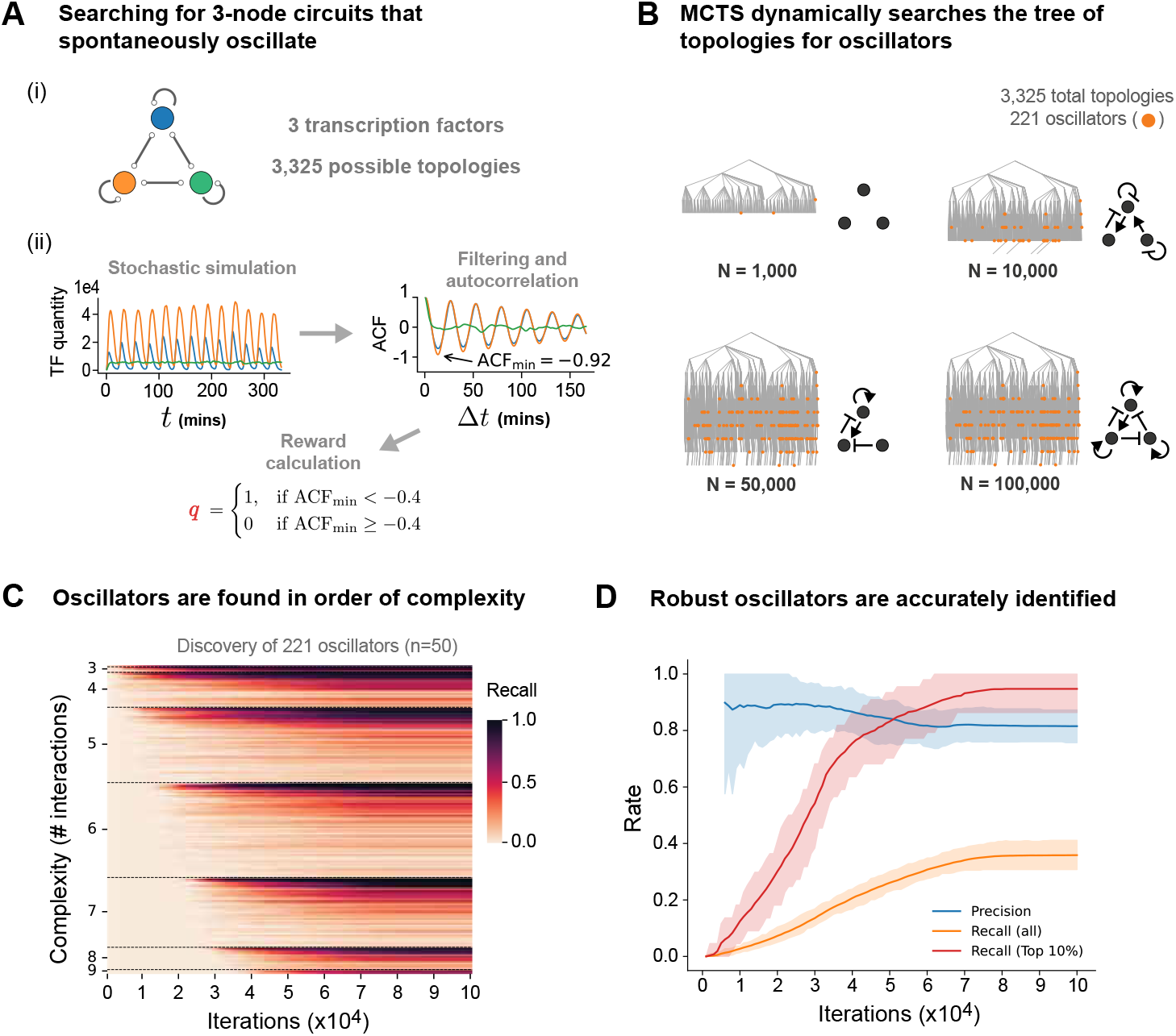
CircuiTree efficiently identifies simple and robust 3-component oscillators. (A) All three-component transcription factor (TF) circuits (3,325 up to symmetry) were enumerated with 10^4^ random parameter sets (i) and evaluated for oscillations (ii) using an autocorrelation-based reward function. (B) A representative MCTS run. With more iterations (*N*), the search graph *T* (represented by a spanning tree for simplicity) expands to encounter more oscillators (orange circles) and improve its best predicted oscillator topology (shown in black). (C) A heatmap showing the average rate of discovery, or recall, for each oscillator (proportion of *n* = 50 replicates). Rows (oscillators) are sorted in order of complexity, or the number of interactions, and oscillators with the same complexity are sorted by descending robustness *Q*. Sparse oscillators are found before more complex ones, with a preference for the most robust candidates. (D) Precision (blue) and recall (orange) of oscillator classification (mean ± 95% CI, *n* = 50). CircuiTree’s recall is particularly high for the 10% most robust oscillators (red), reaching 94.7% after 10^5^ iterations. See also Figures S1, S2, S3, and S4.

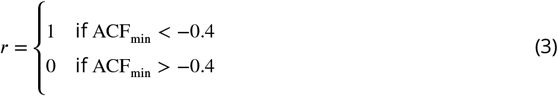

Importantly, the majority of samples classified as oscillatory had periods below 12 minutes (> 55 complete cycles over the simulation), and nearly all had periods below 100 minutes (> 6.6 cycles), indicating that the simulation was long enough to distinguish sustained oscillations from transient dynamics.

#### Algorithm 1

Circuit topology search using MCTS

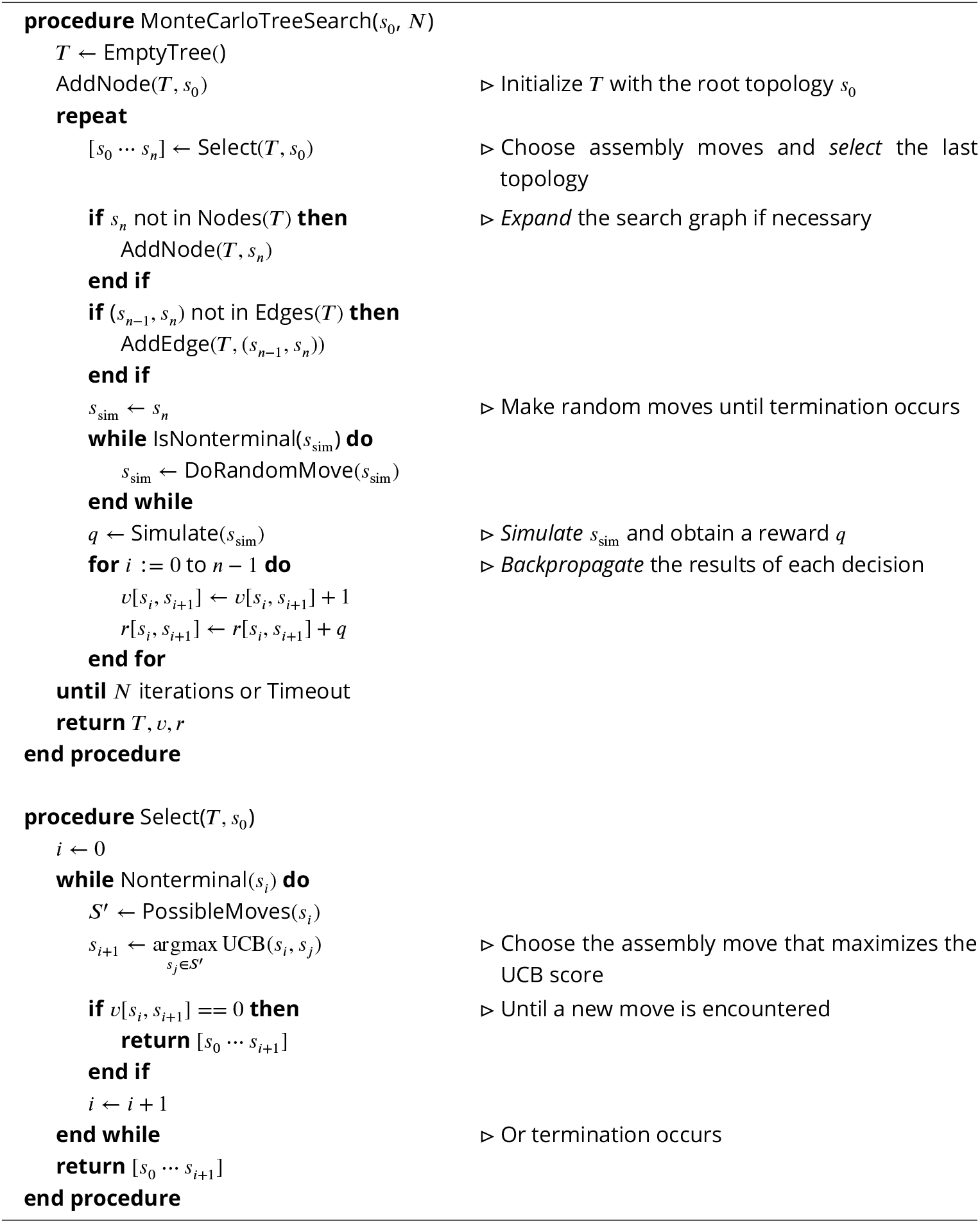

To compare with enumeration, all 3,325 unique, fully connected topologies were simulated with 10^4^ randomly sampled parameter sets and initial conditions. The 10 rate parameters of the model were reduced to 8 dimensionless variables, which were sampled uniformly from a range of values determined based on experimentally measured rates (***Milo et al. (2010***)) and known requirements for oscillations (***Elowitz and Leibler (2000***); ***Novák and Tyson (2008***)). (See Table 2 for variable definitions and sampling ranges and Methods for details of parameter sampling). The robustness to parameter perturbation *Q* was calculated as the fraction of parameter sets that produced oscillation, and topologies with *Q* > 0.01 were considered oscillators. This procedure uncovered 221 oscillator topologies (6.65% of the search space) (see Figures S1B and S1C for a summary of the best topologies). These results generally agree with prior studies of two- and three-node oscillators (***Qiao et al. (2022***); ***Elowitz and Leibler (2000***); ***Novák and Tyson (2008***); ***Woods et al. (2016***); ***Stricker et al. (2008***)). For instance, almost all oscillators contain a repressilator (Rep) loop, activator-inhibitor (A-I) loop, or a combination thereof. Positive autoregulation (PAR), which has been shown to buffer extrinsic and intrinsic noise (***Qiao et al. (2022***)), is also ubiquitous among these oscillators and is required for robust A-I loop oscillations (***Novák and Tyson (2008***); ***Stricker et al. (2008***)). Further discussion of these motifs can be found in the section below. Interestingly, stochastic switching between stable states was occasionally mistaken for low-frequency oscillations (Figure S2), causing some toggle switch circuits to be classified as oscillators (highest *Q* = 0.024, ranked #139).

**Table 2.** Dimensionless variables and limits imposed on random parameter sampling.

| Dimensionless variable | Definition | Sampling limits |
| --- | --- | --- |
| $\kappa_1$ | $\log_{10} [K_{D,1} \cdot 1 \text{ molec}]$ | $[-2, 4]$ |
| $\kappa_2$ | $K_{D,2}/K_{D,1}$ | $[0.00, 0.25]$ |
| $k'_{\text{unbound}}$ | $k_{\text{unbound}} \cdot 1 \text{ sec}$ | $[0, 1]$ |
| $k'_{\text{act}}$ | $k_{\text{act}} \cdot 1 \text{ sec}$ | $[1, 10]$ |
| $r_{\text{inh}}$ | $-\log_{10} [k_{\text{inh}}/k_{\text{act}}]$ | $[0, 5]$ |
| $k'_p$ | $k_p \cdot 1 \text{ molec} \cdot 1 \text{ sec}$ | $[0.015, 0.250]$ |
| $\gamma'_m$ | $\log_{10} [\gamma_m \cdot 1 \text{ molec} \cdot 1 \text{ sec}]$ | $[-3, -1]$ |
| $\gamma'_p$ | $\log_{10} [\gamma_p \cdot 1 \text{ molec} \cdot 1 \text{ sec}]$ | $[-3, -1]$ |

Across all oscillators, oscillation was favored by tight TF-RE binding (*K*_*D*,1_ < 10^3^), low basal transcription (*k*_*m*,unbound_ < 0.3 sec^−1^), strong repression (*k*_*m*,rep_/*k*_*m*,unbound_ < 10^−1^), and a high activated transcription rate (*k*_*m*,act_, monotonic effect) (Figure S3A). Protein and mRNA degradation rates had a non-monotonic effect with a peak at γ_*m*_ ≈ γ_*p*_ ≈ 0.2 molec^−1^ sec^−1^ (Figure S3A), and oscillation period depended strongly on these rates and their ratio (Figures S3B and S3C). However, the best oscillators were exceptionally robust to the choice of parameters. For example, the most robust oscillator (the repressilator with PAR) has a *Q* of 0.767, meaning that for each of the 8 sampled variables, on average, 96.7% of the possible values would permit oscillations (0.967^8^ ≈ 0.767).

### CircuiTree efficiently and systematically discovers three-node oscillators

MCTS masters a game with a limited number of trials by balancing deep sampling of promising regions with broad sampling of under-explored regions. To understand how this strategy performs for circuit topology design, we use CircuiTree to search for 3-node oscillators given *N* = 10^5^ iterations of MCTS (*n* = 50 replicates; see Methods for MCTS implementation specifics). In the average run, the first putative result is discovered after just 0.69 samples/topology (2,280 iterations), and by 19.6 samples/topology (65,360 iterations), a top-5 oscillator has been sampled >100 times. A topology is discovered as a “successful” oscillator if its estimated robustness 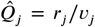 exceeds the threshold value *Q*_thresh_ = 0.01, which was chosen heuristically prior to the search. Figure 2B shows a representative example of how, over sampling time, CircuiTree incrementally builds the search tree from the root state (top to bottom), encounters more oscillators (shown in orange), and improves its prediction for the most robust design (shown as a black circuit diagram). By the end of sampling time, the average run saturates the tree, sampling 99.97% of topologies at least once.

CircuiTree balances efficiency and comprehensiveness by first finding sparse solutions before exploring deeper areas of the tree. In Figure 2C, each row of the heatmap is one of the 221 oscillators, and the rows are sorted first by the circuit’s complexity, or the number of interactions, then by decreasing robustness (*Q*). The color scale indicates the likelihood of discovery, or recall, of each oscillator measured as the proportion of replicates that discovered it. Because sparse topologies require fewer assembly steps, MCTS encounters the sparsest oscillators (such as the repressilator) first, and it discovers increasingly complex solutions over time until oscillators with 9 interactions (the maximum) are found at *N* ≈ 5 × 10^4^. As shown by the line plot in Figure 2D, CircuiTree has a very high recall of 94.7% for the top 10% of oscillators (95% CI: 86.4% − 100.0%) and has a recall of 35.9% (95% CI: 30.8% − 41.5%) for oscillators in general. This is fairly high, considering that it gets an average of 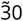 samples/topology, and a majority of oscillators have a *Q* of less than 1/30. Additionally, CircuiTree has a precision of 81.5% (95% CI: 74.1% − 88.3%), indicating a low rate of false positives.

The competing goals of breadth and depth manifest during tree search as distinct temporal phases in which MCTS first explores a broad set of topologies before focusing on a narrow, enriched subset. Among topologies containing a combination of AI and Rep feedback loops, 27.2% (47/173) are oscillators, a very high proportion compared to 6.7% (221/3,325) for the entire design space and 9.3% (146/1,575) and 8.0% (6/75) for the AI or Rep loops alone (Figure S4A). Figure S4B shows how MCTS allocates samples to each of these categories over a total sampling time of 10^5^ and 10^6^ iterations (mean of *n* = 50 and *n* = 12 replicates, respectively). During an initial exploratory phase, samples are taken broadly across categories; however, at *N* ≈ 6 × 10^4^, the AI-Rep combination becomes heavily favored, followed by Rep alone, for the rest of sampling time. This transition can be observed directly from the search history by measuring regret. Defined as

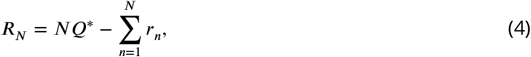

regret is the difference between the expected reward from sampling the best topology (with robustness *Q*^∗^) and the actual reward over *N* iterations. As MCTS moves from exploration into exploitation, the reward rate becomes higher and regret accumulates more slowly (Figures S4C and S4D). Thus, the discovery of an enriched region during search triggers a shift to more focused sampling that can be identified in real-time without *a priori* knowledge about design principles or motifs.

### CircuiTree infers oscillator motifs from search results

In addition to finding individual circuit topologies, an important goal of circuit design is to identify specific structural features, or network motifs, that lead to successful designs (***Milo et al. (2002***); ***Alon (2007***); ***Ma et al. (2009***); ***Cotterell and Sharpe (2010***); ***Shah and Sarkar (2011***); ***Chau et al. (2012***); ***Lim et al. (2013***); ***Schaerli et al. (2014***)). Similarly, in our game-playing paradigm, a motif of the assembly game is a series of moves (an assembly strategy) that leads to a subset of *T* highly enriched with successful design results. Specifically, we define an assembly motif as any successful topology (*Q* > *Q*_thresh_) that is also overrepresented as a sub-pattern within other successful topologies, compared to a set of randomly assembled designs. Using a statistical test for overrepresentation (described in detail in the Methods section and shown schematically in Figure S5A), CircuiTree can mine the results of a tree search for putative assembly motifs.

Looking for 3-node oscillator motifs with ≤ 3 interactions, CircuiTree identifies the same four motifs as enumeration, shown at the top of Figure 3: the repressilator motif (Rep) and the activator-inhibitor (AI) loop with either PAR of the activator (AIPAR), constitutive inhibition of the inhibitor (AICI), or constitutive activation of the activator (AICA). CircuiTree finds these minimal motifs in 100% (Rep), 86% (AIPAR), 94% (AICI), and 62% (AICA) of replicates. To see how these motifs are situated in the overall design space, we plot all 221 oscillators in Figure 3 as a “complexity atlas” (***Cotterell and Sharpe (2010***); ***Schaerli et al. (2014***)), a graph-of-circuits where every oscillator is a node and nodes are organized in layers according to their complexity. Edges connect nodes in adjacent layers if they differ by the addition of a single circuit interaction, analogous to a move in the assembly game. Notably, 216/221 oscillators constitute a large connected component of the graph originating from the four minimal motifs, indicating that almost all 3-node oscillators live in a subset of design space defined by Rep and/or AI motifs. The size of each node in Figure 3 denotes its motif robustness *Q*_motif_, the overall oscillation probability for all circuits containing this motif (in other words, the mean reward once reaching this state in the game). The color of each node reflects its motif discovery rate, measured as the percentage of *n* = 50 replicates that labeled it as a motif.

**Figure 3.**
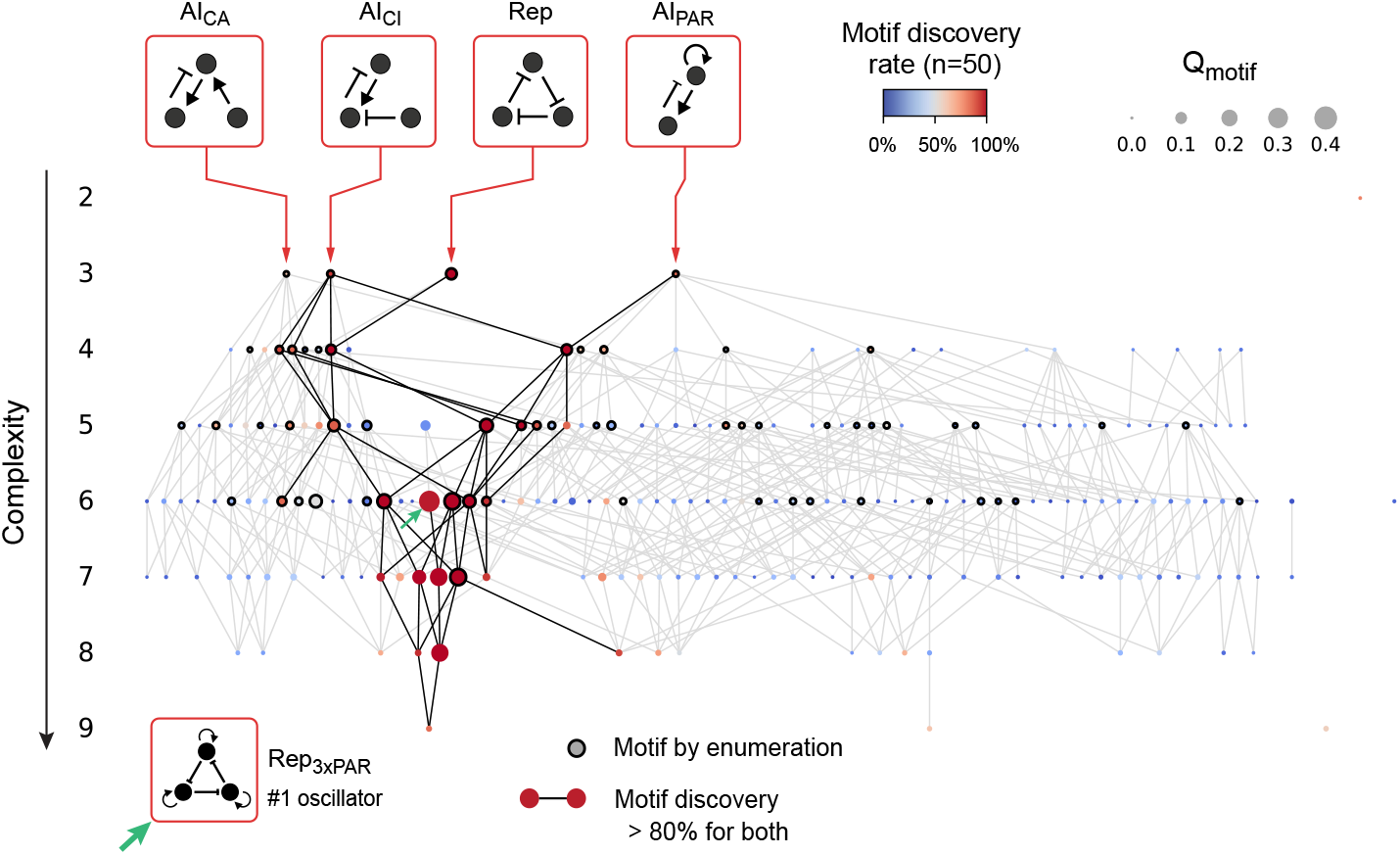
Motifs identified from search results form a cluster of optimal 3-node oscillators. A complexity atlas of oscillators with ≤3 components. Circles are oscillator topologies identified by enumeration, and edges link oscillators that differ by the addition/removal of one interaction. 97.7% of oscillators (216/221) are topologically related to one of the four motifs for 3-node oscillation, shown above the atlas in red boxes. Bold circle borders indicate oscillators found to be motifs based on enumeration. Circle color indicates the rate with which CircuiTree labels each oscillator as an assembly motif. Circle size indicates *Q*_motif_, the average robustness for a circuit completed randomly starting from this state of the assembly game. The correlation between discovery rate and *Q*_motif_ (plotted in Figure S5B) suggests that motifs found by CircuiTree correspond to beneficial game states. The bolded edges, which connect oscillators with a discovery rate > 80%, form a contiguous cluster representing optimal assembly strategies. The most robust oscillator, the repressilator with PAR of all components, is shown on the bottom-left and indicated on the atlas by a green arrow. See also Figures S4 and S5.

Note that these features correlate visually (large circles tend to be red and *vice versa*) and quantitatively, as shown by the scatterplot in Figure S5B, indicating that higher quality motifs are more likely to be found. Additionally, the motifs that are discovered most reliably (discovery rate <80%, indicated by bold edges) form an optimal subset corresponding to the best motifs (the largest nodes). Surprisingly, the most robust oscillator, the repressilator with PAR on all three TFs (Rep+3xPAR, *Q* = 0.767), is discovered at a rate of 98%. Because Rep+2xPAR is a poor oscillator (ranked #187) and Rep+1xPAR does not oscillate at all, Rep+3xPAR cannot be assembled from Rep (or any other intermediate) without breaking oscillations. Nonetheless, its high motif discovery rate suggests that CircuiTree is capable of identifying special design strategies that require a specific combination of moves. Overall, we find that CircuiTree reliably infers the most robust motifs for a given phenotype, even finding obscure but optimal solutions, and the resulting motifs for 3-node oscillation form a single cluster of optimal designs. Note that, in contrast to classic network motifs which compare the frequency of a pattern in successful topologies against the entire set of topologies (or a comparable null model), we compare frequencies between circuit assemblies that were successful during search and random circuit assemblies. Therefore our method returns motifs that are overrepresented *among search results* and thereby accounts for the bias inherent to sampling the space of topologies by assembly rather than by flat enumeration.

### Motif multiplexing allows five-node oscillators to compensate for deletions and single-gene knockdowns

While many studies have elucidated important design principles based on small (1–4 node) network motifs, novel dynamical phenotypes have been noted to emerge in more complex networks. For instance, rather than relying on a single, minimal functional module, circadian oscillators across divergent taxa contain two or more oscillatory feedback loops (***Lee et al. (2000***); ***Cheng et al. (2001***); ***Bell-Pedersen et al. (2005***); ***Pokhilko et al. (2012***)). Could these additional modules make circadian oscillators resistant to genetic perturbations (***Wagner (2005***))? To explore this possibility, we implemented a parallelized version of CircuiTree with pruning and searched for 5-node circuit topologies that oscillate despite a 50% chance of deletion of a random TF (Figures 4A and S6; see methods for details of parallelization). In increasing the circuit size, our principal computational bottleneck was the runtime of the Gillespie algorithm, which scaled linearly with the number of regulatory interactions. Accordingly, to keep the average simulation time under 12 seconds, the 5-node search included only topologies with up to 15 interactions, which represents a search space of approximately 2 billion topologies (see Methods for derivation).

**Figure 4.**
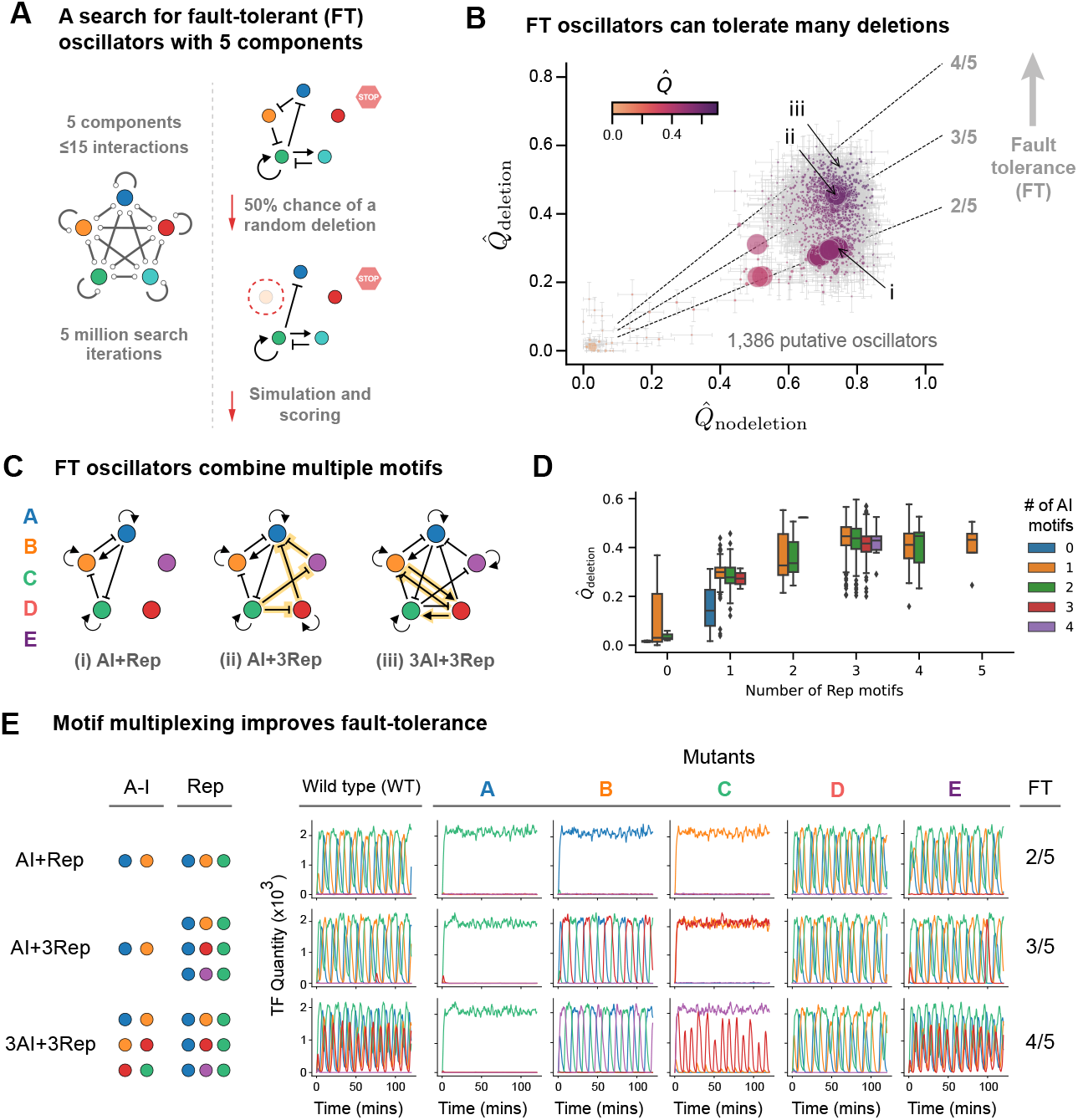
Parallelized CircuiTree scales to large design spaces. (A) A parallelized version of CircuiTree was used to search for five-node oscillators with ≤ 15 interactions (left) that oscillate despite a 50% chance of a single random deletion (right). (B) Search results after 5 ⋅ 10^6^ iterations. Circles are putative oscillators (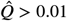, *v*_*i*_ > 100), plotted on axes of robustness based on samples with (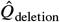) or without a deletion (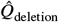). Error bars indicate standard error of the mean. Circle size and color indicate the number of samples *v*_*i*_ and the overall robustness 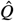, respectively. Dashed lines show different theoretical values of fault-tolerance (FT), or the average number of deletions a circuit can sustain. Examples of circuits with different FT (labeled i-iii) are shown in (C). (D) Box plots of 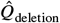, grouped by the number of motifs in each putative oscillator. Multiple motifs, particularly Rep, increase robustness to deletions. (E) Simulated trajectories for the topologies in (C) for all single deletions. The 3AI+3Rep oscillator (FT ≈ 4/5) contains 6 interleaved oscillator motifs. See also Figures S6 and S7.

After 5 million search iterations (less than four days of real time using 1,000 parallel search threads and 300 CPUs), CircuiTree finds 1,386 putative oscillators with 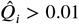 and > 100 samples, the first result arriving in just 1.3 hours. As shown by the heatmap in Figure S7A, when clustered based on their structural similarity (measured as the pairwise graph edit distance between topologies), these topologies separate into sparse and dense clusters (Figure S7B). In Figure 4B, each of these topologies is plotted as a circle based on their observed robustness with or without a random deletion (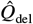 and 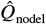, respectively). The overall robustness 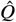 is indicated by the color gradient, and the circle size represents the number of samples. The circuit’s fault-tolerance (FT), the proportion of components that can be deleted without losing oscillation, is estimated as 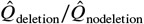, and contour lines for 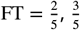, and 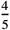 are shown as dashed lines. The putative oscillators included 42 topologies with 2 or 3 nodes, all of which had low robustness to deletions (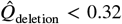). For instance, the second-best oscillator in the 3-node search, the AI+Rep circuit (labeled (i) in Figure 4B and Figure 4C), was found to have a fault tolerance of 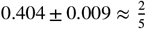, consistent with oscillations that persist after deletion of the two unused TFs (D and E) but attenuating with deletion of any of the active components (A, B, or C).

In contrast, fault-tolerant oscillators (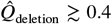) contain many interleaved oscillatory motifs that activate under different deletion scenarios, a design feature we call motif multiplexing. To illustrate this phenomenon, we selected representative circuit topologies that exhibited high robustness both with and without deletion, confirmed their oscillatory behavior using the default parameter set, and identified the combinations of motifs they contained. As shown in Figure 4D, the number of repressilator (Rep) and activator-inhibitor (AI) motifs is a strong predictor of robustness in the presence of deletion, measured as 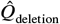, and higher complexity in general is associated with higher 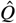 (Figure S7C). To see this design principle, consider the AI+3Rep circuit (labeled (ii) in Figures 4B and 4C), which is an extension of the AI+Rep topology with two additional backup repressilator loops (A-C-D and A-C-E, highlighted in yellow). This circuit has a higher fault tolerance of 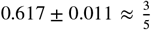 because, while deletion of gene B is fatal for oscillations in the AI+Rep circuit, the A-C-D motif takes over to rescue oscillations in the AI+3Rep circuit (representative simulated trajectories shown in Figure 4E, upper and middle row; see Table 1 for parameter values). The pattern extends to the 3AI+3PAR circuit (labeled (iii) in Figures 4B and 4C, right diagram) which is similar to the AI+3Rep except for the addition of two activator-inhibitor motifs (B-D and D-C, highlighted in yellow). This circuit similarly activates a repressilator motif (A-C-E) upon deletion of gene B. Now, however, oscillations are rescued after deletion of gene C by activating the B-D motif. Consequently, the 3AI+3PAR circuit has a high fault tolerance of 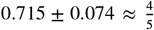 (Figure 4E, bottom row).

During evolution, genomic mutation may lead to a partial reduction in transcription rate rather than a complete knockout. Do multiplexed oscillators maintain their mutational robustness in these conditions? To explore this question, we simulated the 3AI+3Rep circuit under conditions where a single gene is partially knocked down by multiplying its transcription rate by a factor (100 − KD)/100, KD being the percent of knockdown. In Figure 5A, we visualize a representative stochastic simulation of the wild-type (WT) 3AI+3Rep circuit by reducing the system of five TFs (a five-dimensional phase space) to two composite dimensions using principal components analysis and plotting a simulated trajectory on the first and second principal component axes. At each time point, the dominantly expressed TF is indicated by a transparent marker of the same color, and the inset diagram shows the order of TFs activated in the limit cycle (A-B-D-C). As gene A is knocked down from KD=0% to KD=100% (Figure 5B, upper middle panel), the limit cycle gradual drifts in phase space until the trajectory eventually flattens, consistent with the lack of oscillations after deletion of gene A (Figure 4E). During knockdown of gene B (Figure 5B, top right panel), in contrast, the limit cycle drifts before discontinuously jumping to a new limit cycle (the A-E-C repressilator motif) between 75% and 100% KD, as shown in the inset diagram. Similarly, knockdown of gene C induces a gradual drift followed by a jump between 75% and 100% KD to a limit cycle driven by the B-D activator-inhibitor motif. Unlike genes B and C, knockdown of gene D produces no obvious discontinuities as the limit cycle gradually adapts from A-B-D-C to the A-B-C repressilator. Knockdown of gene E produces no appreciable changes in the limit cycle. Overall, the multiplexed oscillator 3AI+3Rep maintains oscillatory behavior during partial knockdown of genes B, C, D, or E and transitions between disparate limit cycles in qualitatively unique ways.

**Figure 5.**
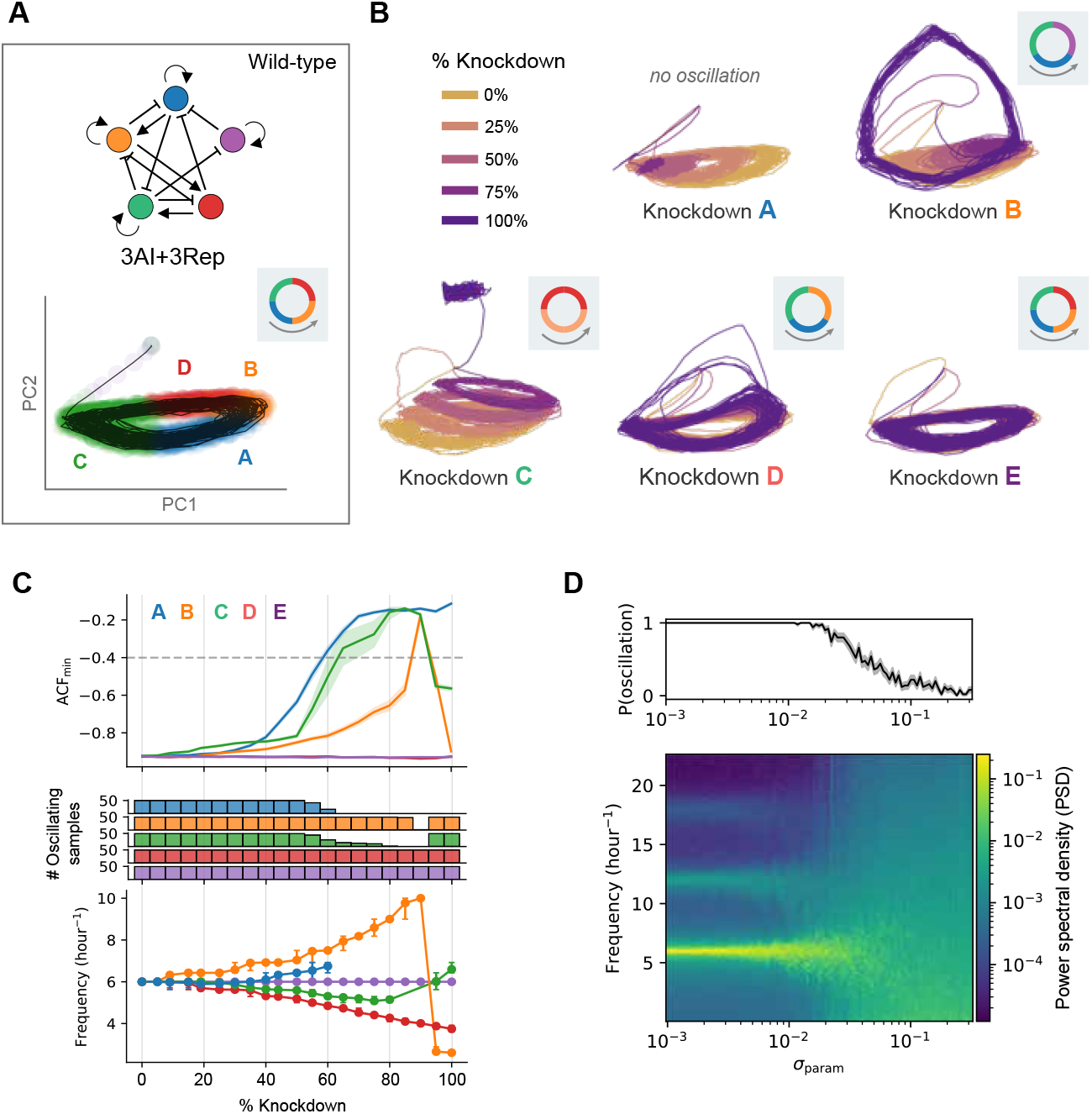
Motif multiplexing makes oscillators resistant to the failure of components. The 3AI+3Rep circuit (A, top) oscillates with different limit cycles after partial knockdowns of different genes. (A, bottom) An exemplary trajectory of 3AI+3Rep is shown on axes of the first two principal components (PCs) of phase space. Transparent circles indicate the dominant species at each time-point. (B) Trajectories under scenarios where transcription rate is reduced by a factor KD. The ordering of species in the limit cycle at KD = 100% is shown by the inset diagram. (C) Oscillation quality and frequency in single-gene knockdowns. Oscillations persist (ACF_min_ < ACF_thresh_) for most knockdowns of genes B, C, D, and E (middle). Oscillation frequency (bottom) is pulled from its WT value in knockdowns of A, B, C, and D. (E) Robustness to parameter variation between TFs. The power spectral density of the trajectory of TF A (bottom, mean) and the overall oscillation rate (top, mean ± SEM) are shown for simulations in which parameters were perturbed by a Gaussian kernel of width σ_param_ (*n* = 50 replicates). A dissipation of fundamental and harmonic frequencies and corresponding loss of oscillations occurs for σ_param_ > 5 ⋅ 10^−2^.

To quantitatively understand how the system responds to knockdowns, single-gene knock-downs were simulated with a range of values of KD with *n* = 50 replicates with different random seeds. The quality and frequency of oscillations were then assessed by computing the ACF_min_ and, if oscillations were detected, the frequency of oscillation. In Figure 5C (upper panel), the ACF_min_ (mean ± 95% confidence interval) is plotted as a function of KD, and the dashed line indicates the value ACF_min_ = −0.4 used as a threshold between oscillatory and non-oscillatory dynamics (below and above the line, respectively). Knockdown of gene A leads to a rise in ACF_min_ and bomes lethal for oscillations above KD = 60%. For gene B, ACF_min_ increases before suddenly crossing the threshold to indicate loss of oscillations between *KD* = 85% and 95%, and for gene C, oscillations disappear in the range KD = 65%*to*90%. For genes D and E, no detrimental effects are observed during knockdown. Thus, oscillations persist in almost all cases (91.9% of samples) when partially knocking down genes B, C, D, or E (Figure 5C, middle panel). Partial knockdowns of A, B, C, and D each have distinct effects on oscillation frequency, a phenomenon called frequency pulling (***Heltberg et al. (2021***)). The bottom panel of Figure 5C shows how, for KD < 60%, the WT resonant frequency of 6.0hr^−1^ is pulled higher when knocking down A or B and lower when knocking down C or D (no change for E). At higher values of KD, knockdowns of B and C discontinuously jump from the drifted frequency to a new resonant frequency (2.6hr^−1^ and 6.6hr^−1^, respectively), while the D knockdown transitions smoothly to its new frequency of 3.8hr^−1^. Thus, the 3AI+3Rep oscillator accommodates most partial knockdowns of genes B, C, D, or E by modulating, yet retaining, oscillatory dynamics.

Up to this point, our modeling has assumed that the rates of reactions such as binding, transcription, translation, and degradation are identical between circuit components. To investigate whether the oscillators we discover are fine-tuned for this symmetry, we perturbed the rate parameters for each TF individually. A Gaussian perturbation kernel with standard deviation σ_param_ was applied to each parameter, scaled to the pre-defined ranges of those parameters (see Table 2 for parameter ranges and Methods for details of the perturbation kernel). Stochastic simulations were performed for a range of values of *σ*_param_ between 1 ⋅ 10^−3^ and 3 ⋅ 10^−2^. To assess the effects of parameter perturbation on oscillation, the power spectral density (PSD) of the dynamics of TF A (mean of *n* = 50 replicates with different random seeds) was calculated for each value of *σ*_param_ and plotted as a heatmap (Figure 5D). For *σ*_param_ < 10^−2^, the PSD shows peaks at the oscillator’s fundamental frequency of 6hr^−1^ and first and second harmonics (12hr^−1^ and 18hr^−1^, respectively). As *σ*_param_ increases further to 5 ⋅ 10^−2^, the fundamental frequency peak gradually diffuses, and the oscillation rate drops from 1.0 to ~ 0.5, as shown in the line plot in Figure 5D (envelope indicates 95% confidence interval). Above *σ*_param_ = 5 ⋅ 10^−2^, the oscillation rate drops to near-zero, and the mean PSD shows no visible peaks. Therefore, this oscillator can tolerate a minimal amount of heterogeneity between TFs, above which it appears somewhat sensitive to asymmetry in at least one parameter of the model.

## Discussion

The size and complexity of natural biological networks exceed our current engineering capabilities by orders of magnitude, underscoring the importance of scalable computational methods for the design of synthetic biological circuits. Inspired by state-of-the-art artificial intelligence (AI), we approach circuit design as a game of step-by-step topology assembly, aiming to achieve a target phenotype in simulation. Our computational platform, CircuiTree, searches the space of possible circuit topologies using the reinforcement learning (RL) algorithm Monte Carlo tree search (MCTS), which balances exploitation of promising circuit assembly moves with exploration of other possibilities (Figure 1). CircuiTree is comprehensive enough to infer motifs for a given phenotype (Figure 3) and efficient enough to search large design spaces fruitfully with limited samples (Figures 4A and 4B). Finally, to demonstrate CircuiTree’s scalability, we use a parallelized implementation of the algorithm to interrogate a space of approximately 2 billion possible five-gene circuits in search of mutationally robust oscillators. Successful designs use a novel strategy we call motif multiplexing. Multiplexed oscillators survive deletion and knockdown mutations by densely interleaving up to six different sub-oscillators (Figures 4C-4E and 5A-5C). Unexpectedly, this multiplexed architecture can transition between distinct limit cycles while mostly avoiding chaos, which commonly arises in coupled oscillators (***Heltberg et al. (2021***)).

Interestingly, eukaryotic circadian clocks across diverse phyla contain multiple sub-oscillators interleaved in a manner that resembles motif multiplexing (***Lee et al. (2000***); ***Cheng et al. (2001***); ***Bell-Pedersen et al. (2005***); ***Pokhilko et al. (2012***)). Our results show that multiplexing confers robustness to genetic perturbations, suggesting that multiplexed circuits may enjoy a selective advantage during evolution. However, several differences between our training scenario and natural evolution limit the conclusions that can be drawn by analogy. After a deletion event, we quantify fitness as the presence of spontaneous and sustained oscillations of any frequency, whereas in nature, specific oscillatory properties, such as a 24-hour period, are subject to selective pressure (***Spoelstra et al. (2016***)). We also subject circuits to whole-gene deletions during training to capture their robustness to worst-case mutations, whereas most naturally occurring mutations produce more gradual changes in gene expression or protein function. Finally, our model does not include entrainment, whereas real circadian clocks synchronize to environmental cycles such as light and temperature. Future work could further interrogate the fault-tolerance of natural circadian clocks and elucidate whether network features such as multiplexing confer evolutionary robustness.

CircuiTree’s combination of generalizability and scalability is unique among computational circuit design tools. Developed for general game-playing and planning problems, MCTS can query very large spaces in search of robust topologies and assembly motifs for any measurable phenotype, without restrictions on the modeling or simulation framework, and without needing to enumerate all possible topologies. This property bridges a gap in computational circuit design where generally, methods for finding single topologies (such as evolution (***François and Hakim (2004***); ***François and Siggia (2008***)), mixed-integer optimization (***Otero-Muras and Banga (2016***)), and recurrent neural networks (***Shen et al. (2021***))) do not generalize easily to design principles and/or require a specific mathematical form, while more comprehensive methods (such as enumeration (***Chau et al. (2012***); ***Schaerli et al. (2014***)) and Bayesian sampling (***Woods et al. (2016***))) require an explicit list of topologies. Unlike these approaches, CircuiTree can efficiently generate many candidate designs, and its tree-search framework allows the user to retrospectively infer design motifs, their prevalence, and their locations in the design space.

While the search space interrogated by CircuiTree in our five-node circuit design task is approximately 2–3 orders of magnitude larger than the largest comprehensive study known to us (***Woods et al. (2016***)), modern RL engines routinely search over games with much larger decision trees. The primary limitations we encountered when scaling further were our simulation paradigm and our adoption of the original MCTS algorithm as a proof-of-concept. While the Gillespie stochastic simulation is exact and realistic, its runtime scales linearly with the number of circuit interactions, limiting us to circuits with up to 15 interactions (~12 seconds/simulation on average). While more work is needed to understand CircuiTree’s scaling behavior, we can gain a rough idea by extrapolating using a power law. Power laws have been shown to describe performance scaling in deep learning models (***Rosenfeld (2021***)), language models (***Kaplan et al. (2020***)), and most recently in RL models (***Hilton et al. (2023***); ***Khatri et al. (2025***)). CircuiTree discovered the first oscillator topology after 2.28 × 10^3^ iterations in the 3-node search (3.325 × 10^3^ topologies) and ~ 8 × 10^5^ iterations in the 5-node search (~ 2 × 10^9^ topologies). This yields a power law of # simulations ~ 65 ⋅ # topologies^0.44^. By extension, a search over 7-node circuits (roughly ~ 3^49^/ 7! ≈ 4.75 × 10^19^ topologies) would stretch the current limits of computation, since it would require 2.7 × 10^10^ simulations, or four weeks of computation on 10,000 CPUs (assuming 1 second/simulation).

However, future methodological advancements in CircuiTree could significantly improve its scaling. Like AlphaZero, CircuiTree incorporates no prior knowledge during training, instead learning purely from simulated outcomes. However, CircuiTree currently implements the original (so-called “vanilla”) MCTS learning model. In contrast, modern RL algorithms based on deep learning have proven far more scalable and drive the current state-of-the-art in game-playing AI (***Browne et al. (2012***); ***Świechowski et al. (2022***)). Value networks and policy networks are key components that allow engines like AlphaZero to search enormous decision spaces. While our largest task searched a space of ~2 billion topologies with 5 million iterations, AlphaZero achieved superhuman performance in chess (~ 10^43^ states) after 44 million self-play games and in Go (10^170^ states) after 140 million (***Silver et al. (2018***)). This problem scale is comparable to a 19-node circuit, which represents a space of ~ 10^172^ possible topologies. Incorporating value and policy networks, together with faster simulation paradigms such as ordinary differential equations or neural ODEs, could substantially extend CircuiTree’s reach to much larger circuits.

More broadly, these results suggest that CircuiTree’s search framework can facilitate the study of network properties that only appear at larger scales, such as the emergent dynamics observed in motif combinations (so-called “hypermotifs”) or the myriad computational capabilities of protein networks (***Adler and Medzhitov (2022***); ***Antebi et al. (2017***); ***Klumpe et al. (2023***); ***Chen et al. (2024***); ***Parres-Gold et al. (2025***)). With improvements in scaling, CircuiTree could facilitate the study and design of more complex phenomena that require larger circuits. In particular, designing cell behavior circuits with multiple cell types is challenging because different cell types harbor distinct intracellular networks linked by an overarching cell-cell signaling network. Envision, for example, an engineered multicellular therapy that treats solid tumors with fewer CAR-T cells by combining them with a targeting cell type and a tissue remodeling cell type, employing a division of labor to maximize tumor penetration and immunogenicity while minimizing adverse cytotoxic effects such as cytokine release syndrome. Similarly, an efficient design paradigm could generate candidate circuits for self-organized morphogenesis, which requires coordinating intracellular regulatory logic with cell-cell signaling, adhesion, and mechanical actuation (***Davies (2017***); ***Toda et al. (2018***); ***Fleischer and Barr (1997***)). In short, CircuiTree’s efficient, scalable automated platform could enable a more systematic understanding of complex biological phenomena and support engineering larger systems, such as multicellular circuits.

## Methods

### Modeling and simulation

#### Stochastic model of a transcription factor network

Unlike phenomenological models of gene regulation that employ Hill functions to describe transcriptional regulation, we model each molecular event explicitly using a system of stochastic reactions describing the synthesis and first-order degradation rates of each mRNA and protein, as well as the sequential binding and unbinding rates of TFs at each of the two response elements (REs) on each promoter. Specifically, for each gene *i* (*i* = 1, …, *K*), the model includes the following reactions, where *m*_*i*_ denotes mRNA, *p*_*i*_ denotes protein (TF), and RE_*i*_ describes the promoter occupancy of gene *i*:

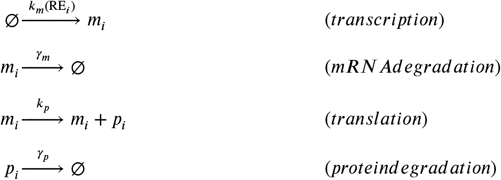

We denote the unoccupied, singly-occupied, and doubly-occupied states of gene *i*’s promoter by 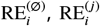, and 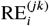 respectively, where *j* and *k* are (possibly identical) TFs. The binding and unbinding reactions of TF *p*_*j*_ to target promoter *i* are:

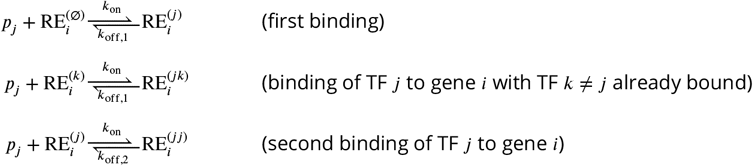

Each interaction between a TF and its target gene is considered either activating or inhibiting in nature. The transcription rate *k*_*m*_ depends on the occupancy of gene *i*’s promoter: *k*_unbound_ (unoccupied), *k*_act_ (one or two activating TFs bound), *k*_inh_ (one or two repressing TFs bound), or *k*_mixed_ = *k*_unbound_ (one activating and one repressing TF bound; see Table 1). The same TF is permitted to act as both an inhibitor and activator of different genes.

#### Sequential binding yields Hill coefficients ≤ 2

In the model, cooperativity arises from the kinetics of sequential binding: when both REs are occupied by the same TF, the second unbinding rate is slower than the first (*k*_off,2_ < *k*_off,1_), stabilizing the doubly-occupied state. Note that in our model, we consider both singly- and doubly-occupied promoters to be transcriptionally active. In the deterministic (continuum) limit, the equilibrium active fraction of promoters as a function of TF copy number *x* is

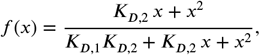

where *K*_*D,i*_ = *k*_off,*i*_/*k*_on_ is the dissociation constant for the *i*-th binding event. The effective Hill coefficient at copy number *x* is defined as *n*_*H*_ (*x*) = *d* ln[*f* /(1 − *f*)]/*d* ln *x*. Computing directly,

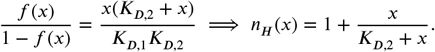

Since *x*/(*K*_*D*,2_ + *x*) ∈ [0, 1) for all *x* ≥ 0, the effective Hill coefficient satisfies *n*_*H*_ ≤ 2 for all TF copy numbers, with equality approached as *x* exceeds *K*_*D*,2_. The same inequality can be shown to hold in the case where only doubly-occupied promoters are transcriptionally active. This result agrees with earlier work by Weiss showing that sequential binding (which he argued is more realistic than multimeric binding) yields relatively weak cooperativity (***Weiss (1997***)).

One practical benefit of sequential binding is a substantial reduction in system complexity, and thus a faster simulation. Consider a circuit diagram of *K* genes (each with mRNA and protein species) with *M* ≤ *K*^2^ regulatory interactions. If (hetero-)dimerization is permitted, the total number of distinct species (counting all mRNAs, TFs, TF dimers, and unique TF-RE complexes) would be 2*K* + *K*^2^ + *M*, and the number of reactions for which propensities must be calculated would be 4*K* + 3*K*^2^ + 3*M* + 3*KM*. In the worst case, the number of propensity calculations scales as *K*^3^, which is quite poor. With sequential binding, the number of distinct species and number of reactions reduce to 2*K* + *M* and 4*K* + 3*M*, respectively, which is much more tractable.

#### Stochastic simulation

Stochastic trajectories were simulated using Gillespie’s exact method (***Gillespie (1977***)) and saved at *n*_*t*_ = 2000 time points at intervals of d*t* = 20 sec. The separation of time-scales between fast binding-unbinding kinetics and the other reactions creates a long simulation time (>2 mins) that cannot be alleviated with, for example, traditional τ-leaping (***Cao et al. (2004***)). Thus, instead of using realistic but impractical binding and unbinding rates, these rates were set to virtual values determined from the equilibrium constant for first-binding *K*_*D*,1_ by the solving the equations

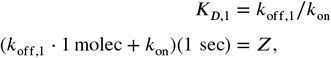

where the value *Z* = 100 was chosen heuristically to be large enough to maintain a separation of time-scales at low quantities. For each protein species, high-frequency noise was filtered from the stochastic signal using a 9-point binomial filter (***Aubury and Luk (1996***)), and the quality of oscillations overall was determined by computing the normalized autocorrelation function for each TF and finding the lowest minimum among TFs (ACF_min_), excluding the bounds. Oscillations were considered present if this quantity, related to the dissipation constant for oscillations (***Otero-Muras and Banga (2016***)), was below the heuristic cutoff ACF_thresh_ = −0.4.

For simulations with partial knockdown of a gene, all transcriptional rate parameters for that gene were multiplied by a coefficient on [0, 1] (for example, KD=80% was achieved using a coefficient of 0.2.), and all species were initialized with zero quantity. For all other simulations, the initial quantity of each TF was selected from a Poisson distribution with a mean of 10 proteins, and all other species were initialized with zero quantity. For perturbation studies, the eight sampled dimensionless variables were converted to values on [0, 1] by normalization to the upper and lower values in Table 2. These values were stored in a *K* × 8 matrix, where each row represented one of the *K* = 5 TFs, and each value was perturbed independently with a Gaussian kernel of standard deviation *σ*_param_, truncated to the range [0, 1] to prevent values outside the reasonable range. The perturbed values were then converted to rate parameters for each TF as outlined above.

### Random sampling

Rate parameters were sampled by drawing 8 dimensionless system variables from a uniform distribution (summarized in Table 2): the log first-binding dissociation constant κ_1_ = log_10_[*K*_*D*,1_ ⋅1 molec] ∈ [−2, 4]; cooperativity ratio κ_2_ = *K*_*D*,2_/*K*_*D*,1_ ∈ [0.00, 0.25]; basal transcription rate 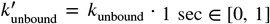; activated transcription 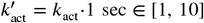; repression ratio *r*_inh_ = − log_10_[*k*_inh_/*k*_act_] ∈ [0, 5]; translation rate 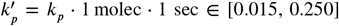; and mRNA and protein degradation rates 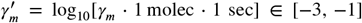 and 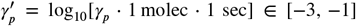. Latin Hypercube sampling was used to draw 10^4^ sets of system variables from this multivariate uniform distribution. For each search (3-node and 5-node), a table was created associating each set of system variables with a unique random seed and initial protein quantities. Each simulation was initialized with a randomly selected row of this table.

### Monte Carlo tree search

For all MCTS runs, the hyperparameter in Equation 2 was set to *c* = 2.00 to encourage exploration (the default value is 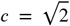). Replicate runs for an experiment were performed using different random seeds, which are used to break ties during the selection phase and draw a random parameter set during the simulation step. For the 3-node search, each reward value *q* was drawn as a Bernoulli trial with a success probability equal to the *Q* value found using enumeration rather than running fresh simulations.

MCTS was parallelized using the lock-free method (***Enzenberger and Müller (2010***)) and implemented with the Python utility celery (https://docs.celeryq.dev/en/stable/index.html). Tree search was parallelized over 1,000 green threads which dispatched simulation jobs to run in parallel on multiple cloud computing instances totaling 300 CPUs. Reward values for each topology-parameter set pair were stored in an in-memory cache to speed up subsequent training runs (epochs). Back-propagation was executed asynchronously with virtual loss, assuming a reward of 0 until the actual reward is returned. To prevent excessive sampling of local optima during the 5-node search, we introduced a form of decision tree pruning we term “node exhaustion”. Once a terminal node (a completed topology) is visited > 10^4^ times, it is considered exhaustively sampled and pruned from the search graph, and a non-terminal node is pruned once all its successors have been pruned. As illustrated in Figure S5, the visits and reward for each in-edge to an exhausted node are subtracted from the parent node. Thus, once a node is marked exhausted, its history is forgotten by its predecessors.

Please see the documentation at https://pranav-bhamidipati.github.io/circuitree/index.html for all additional details of implementation, as well as code tutorials and descriptions of the API.

### Motif identification

Before testing for motifs based on a tree search, we first determine whether each terminated topology *s*_*j*_ discovered during the search is a successful oscillator by comparing its empirical robustness 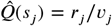 to the predefined threshold *Q*_thresh_ = 0.01. This segregates all topologies into disjoint successful and unsuccessful sets (*X* and *Y*, respectively). We then take samples from the null distribution, drawing *n*_sample_ = 10^5^ samples from the tree of topologies using random assembly, each time starting at the root of the design tree and choosing random actions until a terminal topology is reached (Figure S5A). The procedure is then repeated, this time rejecting the result unless it is a member of the successful set *X* (note that this can be computationally expensive if solutions are sparse). Next, a contingency table is constructed for each successful oscillator *x*_*i*_ ∈ *X* to compare the frequency of observing *x*_*i*_ within samples of the successful subspace against the frequency of observation in the overall space. (see Figure S5A for an illustration of this process). Finally, statistically significant overrepresentation is determined by the χ^2^ independence test with a significance threshold of *p* < *α* = 0.05 after Bonferroni correction. To identify motifs based on the results of enumeration, we conduct the same hypothesis test on each oscillator, except using ground-truth frequencies found by enumeration.

### Estimation of search space size

The size of the 5-node search space was estimated by combinatorial enumeration. Each of the 25 possible interactions (including autoregulatory interactions) between 5 transcription factors can be considered to be an edge on a graph colored with one of three states: absent, activating, or repressing. The number of topologies with exactly *k* non-absent edges is 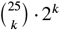, yielding a total of

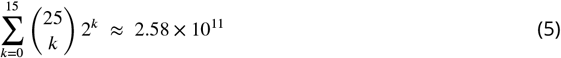

topologies with ≤ 15 interactions. Since the identity of the TF is not considered to matter, topologies that differ only by a relabeling of the TFs are considered equivalent (*A* → *B* → *C* ≅ *B* → *A* → *C*). Topologies that have no internal symmetries are therefore represented 5! = 120 times in Equation 5, once for each permutation of TF labels. If we assume that no topologies exhibit symmetry, we can roughly approximate the number of unique topologies as 2.58 × 10^11^/ 5! ≈ 2.15 × 10^9^. Taking symmetries into account would increase this number; however, restricting this space to connected topologies (i.e. all edges must connect to one another) would decrease this number. Because the net effect of these factors is unknown, this figure should be considered a rough order-of-magnitude estimate.

## Code availability

CircuiTree, written in Python 3.10, is available on GitHub (https://doi.org/10.5281/zenodo.11285522). The code used to run computational experiments, perform analyses, and plot results is available separately at https://doi.org/10.5281/zenodo.11285550.

## Acknowledgments

We thank M. Elowitz, J. Bois, A. Barr, L. Morsut, J. Courte, and J. Paulsson for scientific discussions. We thank J. Bois and G. Manela for advice and assistance on scientific software development. We thank J. Gornet for assistance with distributed computing. We are grateful to I.-M. Strazhnik for assistance with figures and illustrations.

