## Supplementary Figures for "Designing biochemical circuits with tree search"

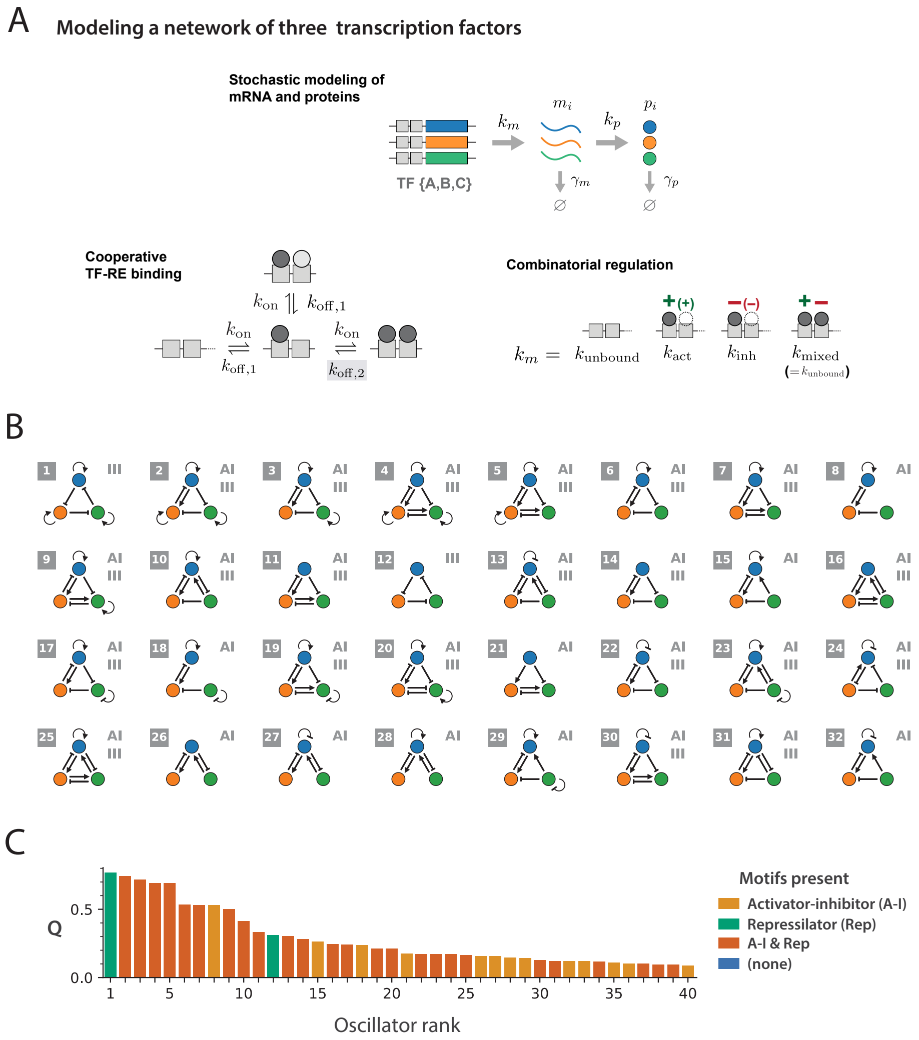


**Figure** S1: **Modeling and enumeration of 3-component oscillators.** (A) Stochastic modeling of a network of 3 transcription factors. See Table 1 for explanations and default values for reaction rate parameters. (B) The top-32 most robust oscillators (highest $Q$) based on enumeration and exhaustive simulation ($10^4$ parameter sets). "AI" and "III" refer to the presence of at least one activator-inhibitor or repressilator motif, respectively. (C) Robustness of the top-40 most robust oscillators. Bars are colored based on the presence or absence of an AI and/or repressilator (Rep) motif. See also Figure 2 and Table 1.

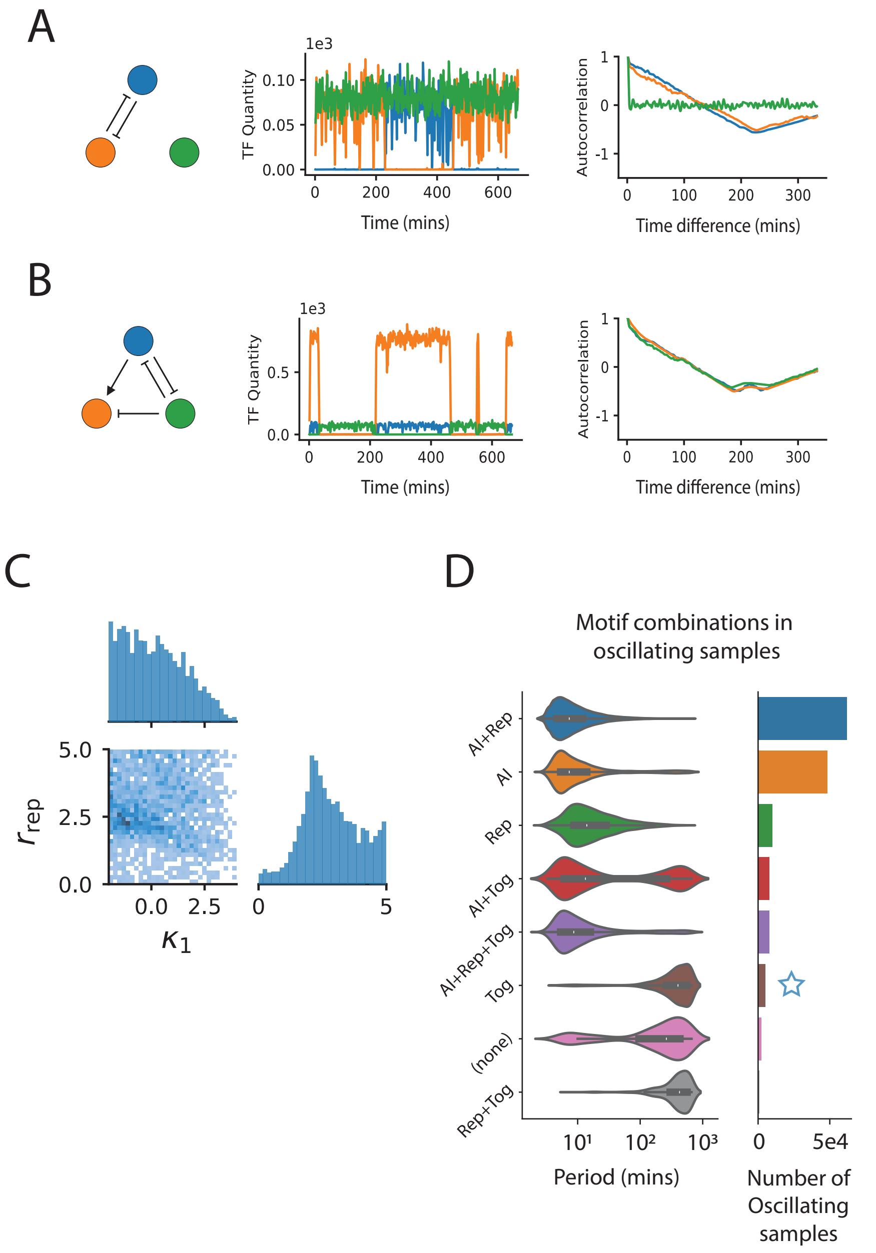


**Figure S2: Strong TF-RE binding and moderate repression enable low-frequency toggle switch "oscillations".**

**Figure** S2: (A-B) Circuit topology, representative stochastic trajectory, and autocorrelation for two toggle switch topologies that were classified as oscillators, the basic toggle switch (A), ranked #148, and an amplified toggle switch (B), ranked #139. Occasional switching between stable states on a timescale comparable with the total simulation time is indistinguishable from low-frequency oscillation. (C) For oscillators containing a toggle switch and lacking either the Rep or AI oscillatory motifs, parameter sets resulting in oscillation generally showed strong TF-RE binding (low $\kappa_1$) and moderately strong repression ($r_{\text{rep}} \approx 2.5$). The latter may be a "sweet spot" that enables a persistence time on the order of the total simulation time. (D) Left: Violin plots of oscillation period for oscillators with different combinations of AI, Rep, and toggle switch (Tog) motifs. Oscillators with a toggle switch appear to have much longer periods, on the order of many hours. Right: Bar plot of the total number of oscillating samples shown in the violin plots. Samples used to generate (C) are denoted with a blue star. See also Figure 2 and Tables 1 and 2.

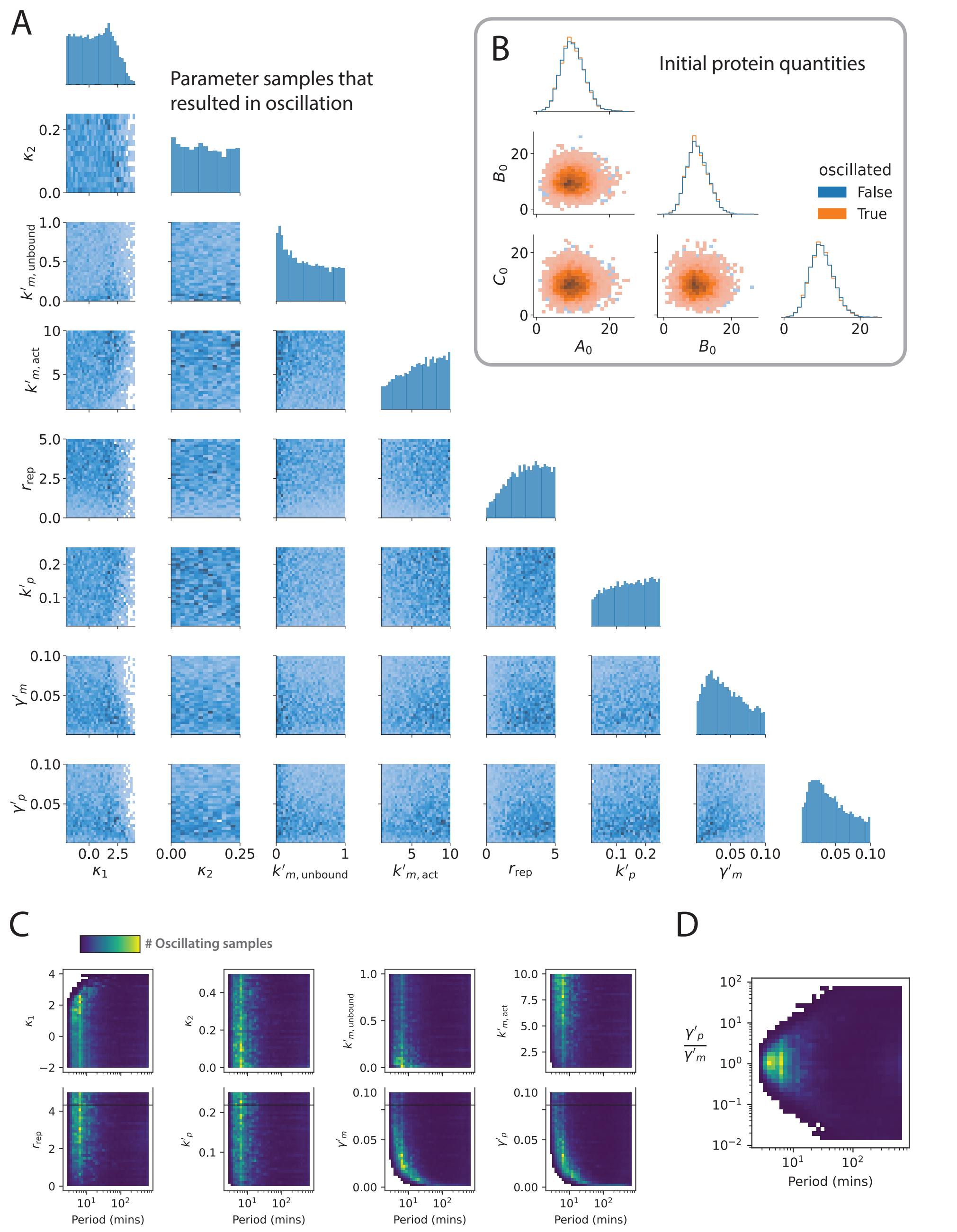


**Figure S3: Dependence of oscillation likelihood and period on sampled variables.**

**Figure** S3: (A) A corner plot of all samples taken during 3-node enumeration that resulted in oscillation, across all topologies. The histograms on the diagonal show the marginal distribution for each sampled variable, and heatmaps in the lower triangular of the grid show every pairwise dependency of these parameters. Weak binding (high $\kappa_1$) and weak repression $lowr_{\text{rep}}$ are the only visibly prohibitive parameter regimes for oscillation. Oscillations are favored by parameter sets with low basal expression (low $k'_{m,\text{unbound}}$), strong activation (high $k'_{m,\text{act}}$), and comparable mRNA and protein degradation rates ($\gamma'_m \approx \gamma'_p \approx 0.02$). (B) No dependence on initial protein quantities was noted. (C) 2D density plots showing the dependence of oscillation period on the variables. Only $\gamma'_m$ and $\gamma'_p$ seem anticorrelated with period, otherwise no dependence is noted. (D) Oscillation period, unlike oscillation likelihood, does not seem to depend on $\gamma'_p/\gamma'_m$. See also FIgure 2 and Tables 1 and 2.

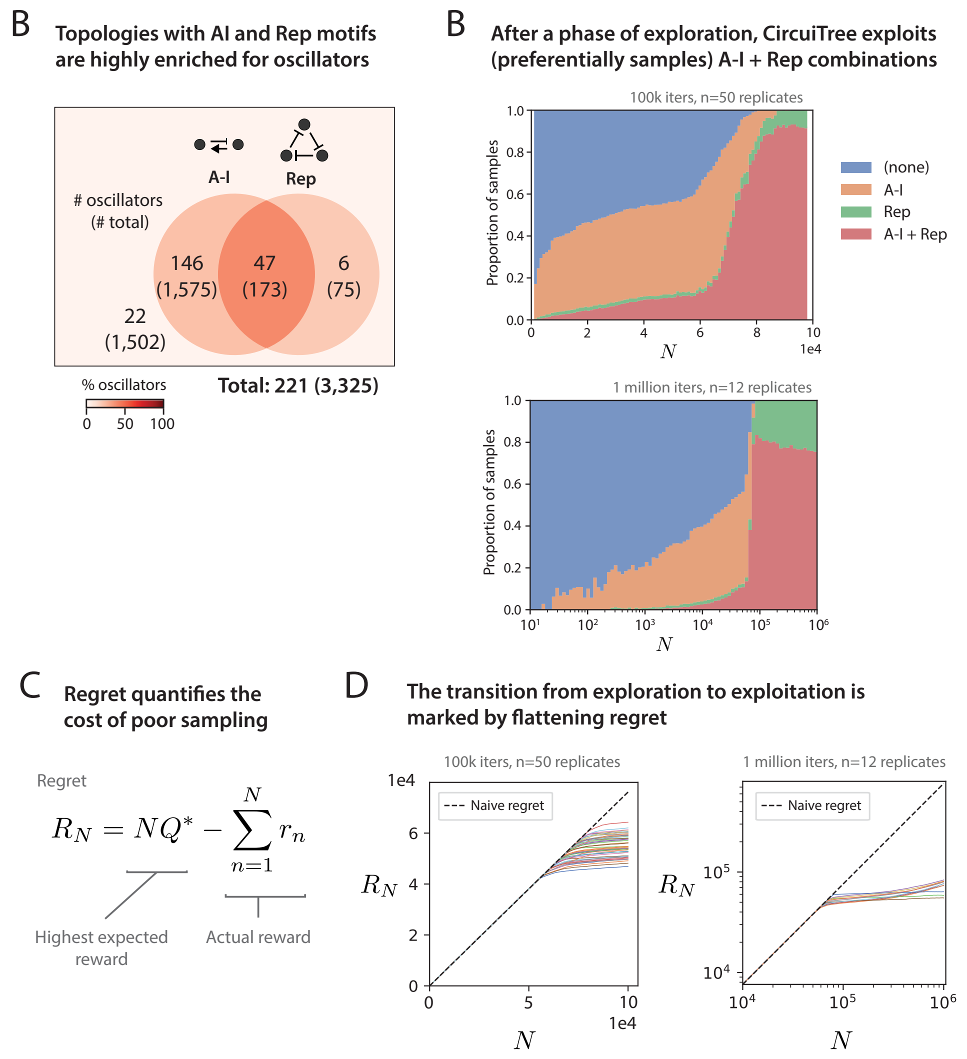


$$R_N = NQ^* - \sum_{n=1}^{N} r_n$$

**Figure S4: After a period of exploration, CircuiTree reduces sampling regret by exploiting motif combinations.** (A) Venn diagram of 3-node oscillators discovered by enumeration, grouped by presence or absence of AI or Rep motifs. For each category, the number of oscillators (number of total topologies) is shown. The intensity of background color denotes the percentage of topologies in that category that are oscillators. Notably, 27.2% (47/173) of topologies with the AI-Rep combination are oscillators.

**Figure** S4: (B) The proportion of samples allocated to each motif combination for MCTS runs lasting $10^5$ iterations (top; mean of $n = 50$ replicates; linear x-axis) and $10^6$ iterations (bottom; mean of $n = 12$ replicates; logarithmic x-axis). There is a gradual shift towards sampling AI and AI-rep combinations before, at around $6 \times 10^4$ iterations, the AI-Rep motif combinations (red) and, to a lesser extent, the Rep-only category (green) are suddenly and dramatically exploited. The effect on the accumulation of reward can be seen by calculating regret. Defined in (C), regret is the opportunity cost accrued by sampling sub-optimal topologies ($Q < Q^*$). (D) At the transition between the initial phase and the exploitation phase (shown on linear axes on the left and log-log axes on the right), regret flattens because exploitation of AI-Rep combinations has increased the rate of reward. See also Figures 2 and 3.

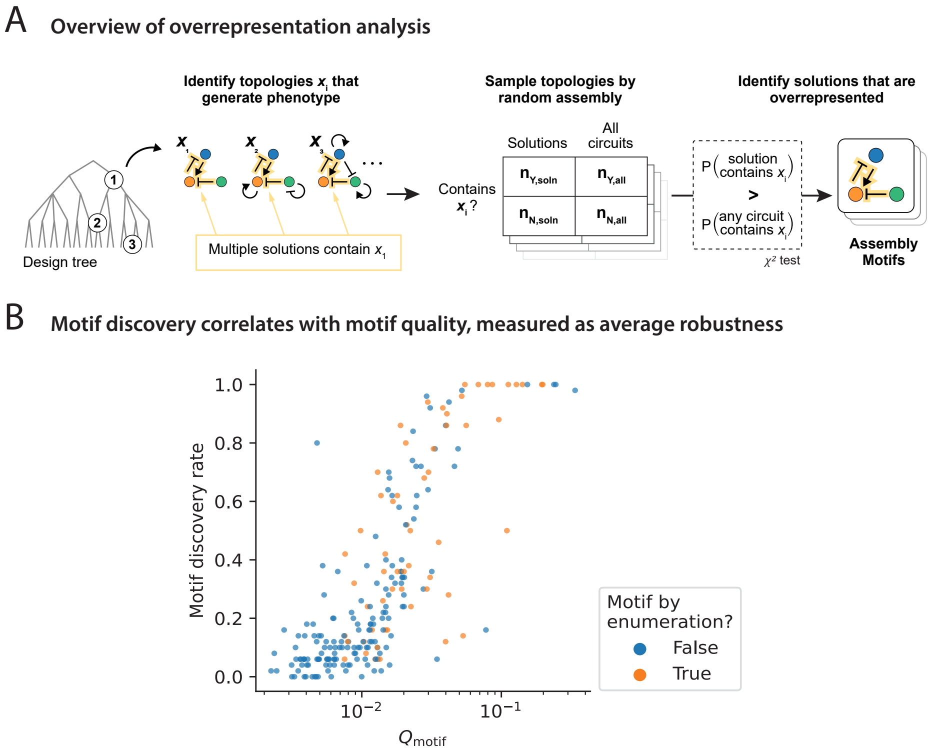


**Figure** S5: **Overrepresentation analysis infers assembly motifs by random sampling of the search graph.** (A) A flowchart of overrepresentation analysis, described in detail in the main Methods section. The goal is to find patterns that are overrepresented relative to the overall design space without needing to explicitly enumerate that space. This is achieved using a random sampling scheme. Unlike traditional design motifs, which are found by comparing to a flat, enumerated null distribution, these assembly motifs are (virtually by definition) good assembly strategies. This is demonstrated by the scatterplot in (B) showing the relationship between $Q_{\text{motif}}$ and motif discovery by CircuiTree. $Q_{\text{motif}}$ can be conceptualized as the average win probability from a given assembly state if taking random actions. Motifs with a high discovery rate by CircuiTree are likely to have high $Q_{\text{motif}}$ and *vice versa*. Meanwhile, motifs found by traditional enumeration (orange circles), while overrepresented among solutions, are not necessarily beneficial moves in the assembly game. See also Figure 3.

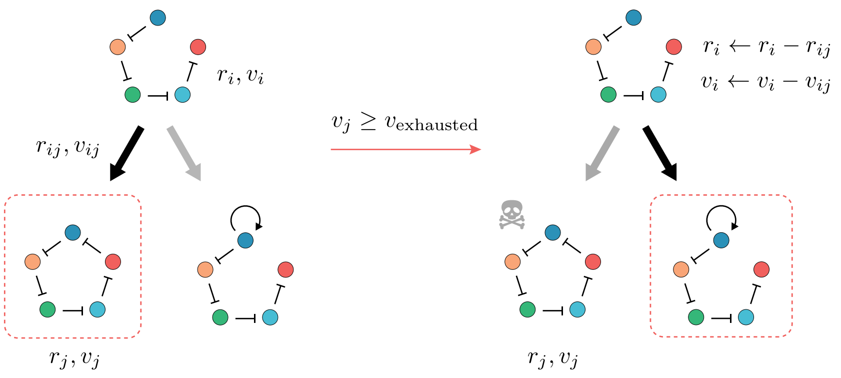


**Figure** S6: **States are pruned based on sampling depth to avoid over-sampling local optima.** A schematic of state pruning implemented in the parallel version of CircuiTree during the 5-node search. While asymptotically comprehensive, MCTS may perseverate for many iterations in a local optimum of design space. To gently discourage this over-focusing, we implement a pruning step during selection in which a selected state that has been visited more than $v_{\text{exhausted}} = 10^4$ times (the boxed circuit on the left) is marked as "exhausted" (in the sense of exhaustively sampled), denoted by the skull and crossbones. An exhausted state $s_j$ can no longer be selected in subsequent iterations, and each of its parent states $s_i$ (each predecessor of $s_j$ in $T$) is made to "forget" the sampling history of $s_j$ by subtracting the visits and rewards of $s_{ij}$ from the totals of the parent $s_i$.

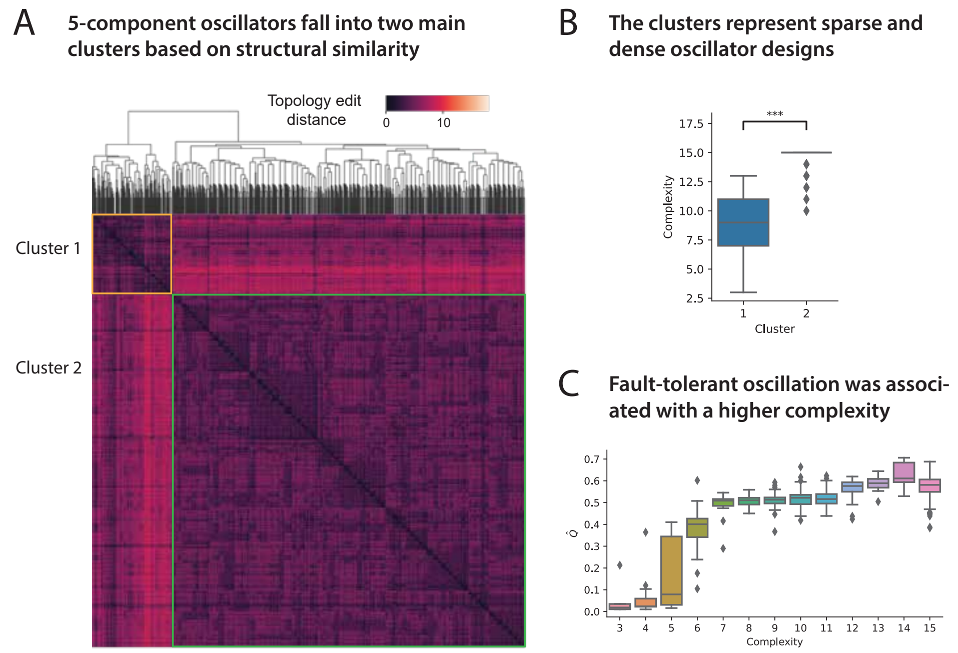


**Figure** S7: **Five-node fault-tolerant oscillators cluster based on topological complexity.** (A) A clustered heatmap of the 1,386 putative oscillators discovered during the search for 5-node fault-tolerant oscillators. Pairwise distance is computed as the graph edit distance between topologies (the number of edges that must be added/deleted to make two topologies equivalent). Putative FT oscillators fall into two structurally similar clusters, cluster 1 (outlined in orange) and cluster 2 (outlined in blue). (B) Box plots of topological complexity, grouped by cluster. Cluster 1 contains relatively sparse topologies, while the majority of topologies in cluster 2 have 15 interactions, the maximum allowed during search. $^{***}p < 0.001$ by Mann-Whitney U-test. (C) Boxplots of overall robustness $\hat{Q}$ grouped by complexity. Regardless of cluster, higher complexity is associated with higher $\hat{Q}$. Specifically, there is a visible jump in robustness at 7 interactions and perhaps again at 14 interactions.
